# Identification of multiple transcription factor genes potentially involved in the development of electrosensory versus mechanosensory lateral line organs

**DOI:** 10.1101/2023.04.14.536701

**Authors:** Martin Minařík, Melinda S. Modrell, J. Andrew Gillis, Alexander S. Campbell, Isobel Fuller, Rachel Lyne, Gos Micklem, David Gela, Martin Pšenička, Clare V. H. Baker

## Abstract

In electroreceptive jawed vertebrates, embryonic lateral line placodes give rise to electrosensory ampullary organs as well as mechanosensory neuromasts. Previous reports of shared gene expression suggest that conserved mechanisms underlie electroreceptor and mechanosensory hair cell development and that electroreceptors evolved as a transcriptionally related ’sister cell type’ to hair cells. We previously identified only one transcription factor gene, *Neurod4*, as ampullary organ-restricted in the developing lateral line system of a chondrostean ray-finned fish, the Mississippi paddlefish (*Polyodon spathula*). The other 16 transcription factor genes we previously validated in paddlefish were expressed in both ampullary organs and neuromasts. Here, we used our published lateral line organ-enriched gene-set (arising from differential bulk RNA-seq in late-larval paddlefish), together with a candidate gene approach, to identify 23 transcription factor genes expressed in the developing lateral line system of a more experimentally tractable chondrostean, the sterlet (*Acipenser ruthenus*, a small sturgeon), and/or that of paddlefish. Twelve are expressed in both ampullary organs and neuromasts, consistent with conservation of molecular mechanisms. Six are electrosensory-restricted on the head *(Irx5*, *Insm1*, *Sp5*, *Satb2*, *MafA* and *Rorc*), and five are the first-reported mechanosensory-restricted transcription factor genes (*Foxg1*, *Sox8*, *Isl1*, *Hmx2* and *Rorb*). However, as previously reported, *Sox8* is expressed in ampullary organs as well as neuromasts in a shark (*Scyliorhinus canicula*), suggesting the existence of lineage-specific differences between cartilaginous and ray-finned fishes. Overall, our results support the hypothesis that ampullary organs and neuromasts develop via largely conserved transcriptional mechanisms, and identify multiple transcription factors potentially involved in the formation of electrosensory versus mechanosensory lateral line organs.

## Introduction

In jawed anamniotes, mechanosensory hair cells are found in the inner ear, in the spiracular organ associated with the first pharyngeal cleft (lost in amphibians, bichirs and teleosts; see Norris and Hughes, 1920; von Bartheld and Giannessi, 2011), and in lateral line neuromasts: small sense organs distributed in lines over the head and trunk, which respond to local water movement (see e.g., Mogdans, 2019; Webb, 2021). In electroreceptive species (e.g., cartilaginous fishes, ray-finned fishes including the chondrostean paddlefishes and sturgeons, and urodele amphibians like the axolotl), the lateral line system includes electrosensory ampullary organs containing supporting cells and electroreceptors that detect weak electric fields, primarily for hunting or avoiding predators (see e.g., Crampton, 2019; Leitch and Julius, 2019; Chagnaud et al., 2021).

Like hair cells, electroreceptors have an apical primary cilium (lost during maturation in cochlear hair cells; Lu and Sipe, 2016) and basolateral ribbon synapses with afferent nerve terminals (Jørgensen, 2005; Baker, 2019). However, electroreceptors lack the apical hair bundle (staircase array of microvilli) where mechanoelectrical transduction occurs (Ó Maoiléidigh and Ricci, 2019; Caprara and Peng, 2022). The main anamniote developmental models - the teleost zebrafish and the frog *Xenopus* - lack ampullary organs: electroreception was lost in the ray-finned bony fish radiation leading to teleosts, and in the lobe-finned bony amphibian lineage leading to frogs (Baker et al., 2013; Baker, 2019; Crampton, 2019). (Physiologically distinct lateral line electroreceptors evolved independently in some teleost lineages; see e.g., Baker et al., 2013; Baker, 2019; Crampton, 2019.)

Neuromasts and ampullary organs, together with the neurons in lateral line ganglia that provide their afferent innervation, develop from a series of cranial lateral line placodes (Northcutt, 1997; Baker et al., 2013; Piotrowski and Baker, 2014). These either elongate to form sensory ridges that fragment, with a line of neuromasts forming first in the ridge’s centre and ampullary organs (if present) forming later on the ridge’s flanks (Northcutt, 1997; Piotrowski and Baker, 2014). Alternatively, as in the ray-finned bony teleost zebrafish, lateral line primordia migrate as cell collectives, depositing neuromasts in their wake (Piotrowski and Baker, 2014). The lateral line placode origin of ampullary organs was first shown by grafting and ablation in a lobe-finned bony tetrapod, the axolotl (Northcutt et al., 1995). Our DiI-labelling studies in a chondrostean ray-finned fish (Mississippi paddlefish, *Polyodon spathula*) (Modrell et al., 2011a) and a cartilaginous fish (little skate, *Leucoraja erinacea*) (Gillis et al., 2012) showed that individual elongating lateral line placodes form ampullary organs and neuromasts across all non-teleost jawed vertebrates (reviewed in Baker et al., 2013).

What molecular mechanisms underlie the formation of ampullary organs versus neuromasts within the same lateral line sensory ridge? We have identified a range of ampullary organ-expressed genes in different electroreceptive vertebrates using a candidate gene approach (O’Neill et al., 2007; Modrell et al., 2011a; Modrell et al., 2011b; Gillis et al., 2012; Modrell and Baker, 2012; Modrell et al., 2017a; Modrell et al., 2017b). More recently, we took an unbiased differential RNA-seq approach, comparing the transcriptome of late-larval paddlefish gill-flaps (covered in ampullary organs, plus some neuromasts) versus fins (no lateral line organs) (Modrell et al., 2017a). The resultant lateral line organ-enriched dataset of around 500 genes (Modrell et al., 2017a) is not exhaustive: it includes most, but not all, of the genes identified in paddlefish ampullary organs via the candidate gene approach (Modrell et al., 2011a; Modrell et al., 2011b; Modrell et al., 2017a; Modrell et al., 2017b). *In situ* hybridization for selected candidate genes from this dataset suggested that electroreceptors and hair cells are closely related, e.g., late-larval ampullary organs express genes encoding proteins essential for neurotransmission specifically at hair-cell (but not photoreceptor) ribbon synapses in the basolateral cell membrane (Pangrsic et al., 2018; Moser et al., 2020), such as vGlut3, otoferlin and Ca_v_1.3 (Modrell et al., 2017a). Ca_v_1.3 has also been identified as the electrosensitive voltage-gated Ca^2+^ channel in the apical electroreceptor membrane (Bennett and Obara, 1986; Bodznick and Montgomery, 2005) in shark and skate species (i.e., in cartilaginous fishes) (Bellono et al., 2017; Bellono et al., 2018).

Developing ampullary organs also express key ’hair cell’ transcription factor genes including *Atoh1*, *Pou4f3* (*Brn3c*) and *Six1* (Modrell et al., 2011a; Butts et al., 2014; Modrell et al., 2017a). These are critical for hair cell formation and, in combination with *Gfi1*, can drive an immature ’hair cell-like’ fate in mouse embryonic stem cells, adult cochlear supporting cells and fibroblasts (see Roccio et al., 2020; Chen et al., 2021; Iyer and Groves, 2021; Iyer et al., 2022). Co-expression of *Atoh1*, *Pou4f3* and *Gfi1* is sufficient to drive a more mature hair cell fate in postnatal cochlear supporting cells (Chen et al., 2021). The expression of *Atoh1*, *Pou4f3* and *Six1* in developing ampullary organs, as well as neuromasts, suggests that molecular mechanisms underlying electroreceptor development are likely to be highly conserved with those underlying hair cell formation (Modrell et al., 2011a; Modrell et al., 2017a). Nevertheless, hair cells and electroreceptors are morphologically and functionally distinct (Jørgensen, 2005; Baker, 2019). Validation of multiple candidate genes from the late-larval paddlefish lateral line organ-enriched dataset had identified only a handful of genes with expression in developing ampullary organs but not neuromasts (Modrell et al., 2017a). These were the basic helix-loop-helix (bHLH) transcription factor gene *Neurod4*, plus two voltage-gated potassium channel subunit genes (*Kcna5*, encoding K_v_1.5, and *Kcnab3*, encoding the accessory subunit K_v_ý3) and a calcium-chelating beta-parvalbumin, all presumably involved in electroreceptor function (Modrell et al., 2017a).

In recent years, another chondrostean fish, the sterlet (a sturgeon, *Acipenser ruthenus*), has been developed as an experimentally tractable non-teleost model (see e.g., Saito and Psenicka, 2015; Chen et al., 2018; Baloch et al., 2019; Stundl et al., 2020; Du et al., 2020; Stundl et al., 2022; Stundl et al., 2023). In contrast to the limited Mississippi paddlefish spawning season, many hundreds of sterlet embryos are available each week for up to several months in fully equipped laboratory research facilities. We have therefore turned to the sterlet as a tractable model for studying the molecular control of lateral line hair cell and electroreceptor development. In the current study, we describe lateral line hair cell and electroreceptor differentiation in the sterlet, and the expression in sterlet and/or paddlefish of almost all of the remaining transcription factor genes from the paddlefish late-larval lateral line organ-enriched dataset (Modrell et al., 2017a), plus a few additional candidates. We report expression within the developing lateral line system of 23 novel transcription factor genes. Twelve - including the key ’hair cell’ transcription factor gene *Gfi1* - were expressed in both ampullary organs and neuromasts, supporting conserved molecular mechanisms. Six transcription factor genes proved to be electrosensory-restricted, while five represent the first-reported mechanosensory lateral line-restricted transcription factors. These eleven genes, plus ampullary organ-restricted *Neurod4* (Modrell et al., 2017a), are good candidates to be involved in the development of electrosensory versus mechanosensory lateral line organs.

## Results

### Characterizing lateral line hair cell and electroreceptor development in sterlet

In order to use the sterlet as a more experimentally tractable model for lateral line organ development than the Mississippi paddlefish (Modrell et al., 2011b; Modrell et al., 2011a; Modrell et al., 2017b; Modrell et al., 2017a), we first characterized the timing and distribution of lateral line hair cell and electroreceptor differentiation (staging according to Dettlaff et al., 1993). The development of lateral line placodes, neuromasts and ampullary organs had previously been described in the shovelnose sturgeon, *Scaphirhynchus platorynchus* (Gibbs and Northcutt, 2004a); ampullary organ formation had also been described in a sturgeon in the same genus as the sterlet, the Adriatic sturgeon, *A. naccari* (Camacho et al., 2007). In *S. platorynchus*, all lateral line placodes are present at stage 30 and have started to elongate into sensory ridges (Gibbs and Northcutt, 2004a). By the time of hatching at stage 36, neuromast primordia are present within the sensory ridges at the centre of all the lateral line placodes but only a few mature neuromasts have formed, specifically in the otic lateral line (Gibbs and Northcutt, 2004a). At stage 41, roughly midway between hatching (stage 36) and the onset of independent feeding (stage 45), ampullary organ primordia are present in the lateral zones of the anterodorsal, anteroventral, otic and supratemporal sensory ridges (Gibbs and Northcutt, 2004a). At stage 45, mature ampullary organs are present in the infraorbital, otic and posterior preopercular fields and ampullary organs continue to develop over the next three weeks (Gibbs and Northcutt, 2004a). By stage 45 in *A. naccari*, in contrast, ampullary organs are ultrastructurally mature at almost all sites and thought to be functional (Camacho et al., 2007). Camacho et al. (2007) speculate that the difference is due to *S. platorynchus* reaching stage 45 on day 4 post-hatching, leaving less time for completion of ampullary organ development than in *A. naccari*, which reaches stage 45 on day 9 post-hatching.

To examine the formation and distribution of developing neuromasts and ampullary organs in sterlet, we focused on the stages between stage 35, i.e., the stage before hatching (stage 36), up to the onset of independent feeding at stage 45, which is reached in the sterlet at day 8 post-hatching (14 days post-fertilization, dpf) (Zeiske et al., 2003). Figure 1 shows a temporal overview of the expression by *in situ* hybridization (ISH) in sterlet of *Eya4*, an otic and lateral line primordium marker that is eventually restricted to differentiated hair cells and electroreceptors (Modrell and Baker, 2012; Baker et al., 2013) (Figure 1A-D) and *Sox2*, which is also expressed by lateral line primordia and is eventually restricted to supporting cells in both ampullary organs and neuromasts, with stronger expression in neuromasts (Modrell et al., 2017a) (Figure 1E-H). (*Sox2* is also expressed in taste buds on the barbels and around the mouth; Figure 1E-H.) As a marker for differentiated hair cells and electroreceptors, we used *Cacna1d*, which encodes the pore-forming alpha subunit of Ca_v_1.3 and is expressed in hair cells and electroreceptors across jawed vertebrates (Modrell et al., 2017a; Bellono et al., 2017; Bellono et al., 2018) (Figure 1I-L; also see Supplementary Figure S1A-H). To identify electroreceptors specifically, we cloned the two voltage-gated potassium channel subunit genes that we identified as electroreceptor-specific in paddlefish (Modrell et al., 2017a): *Kcnab3*, encoding the accessory subunit K_v_ý3 (Figure 1M-P; also see Supplementary Figure S1I-P) and *Kcna5*, encoding K_v_1.5, which shows the same expression pattern as *Kcnab3* (data not shown.) The earliest sign of neuromast hair cell differentiation was at stage 35 (Figure 1I; Supplementary Figure S1A,B) with increasing numbers at all subsequent stages (Figure 1J-L; Supplementary Figure S1C-H). Differentiated electroreceptors were not seen until stages 40-41, in some ampullary organ fields (Figure 1I-K,M-O; Supplementary Figure S1I-N). By stage 45 (the onset of independent feeding), all cranial neuromast lines and fields of ampullary organs with differentiated electroreceptors could be identified (Figure 1D,H,L,P; Supplementary Figure S1G,H,O,P). A schematic summary is shown in Figure 1Q-T.

**Figure 1.**
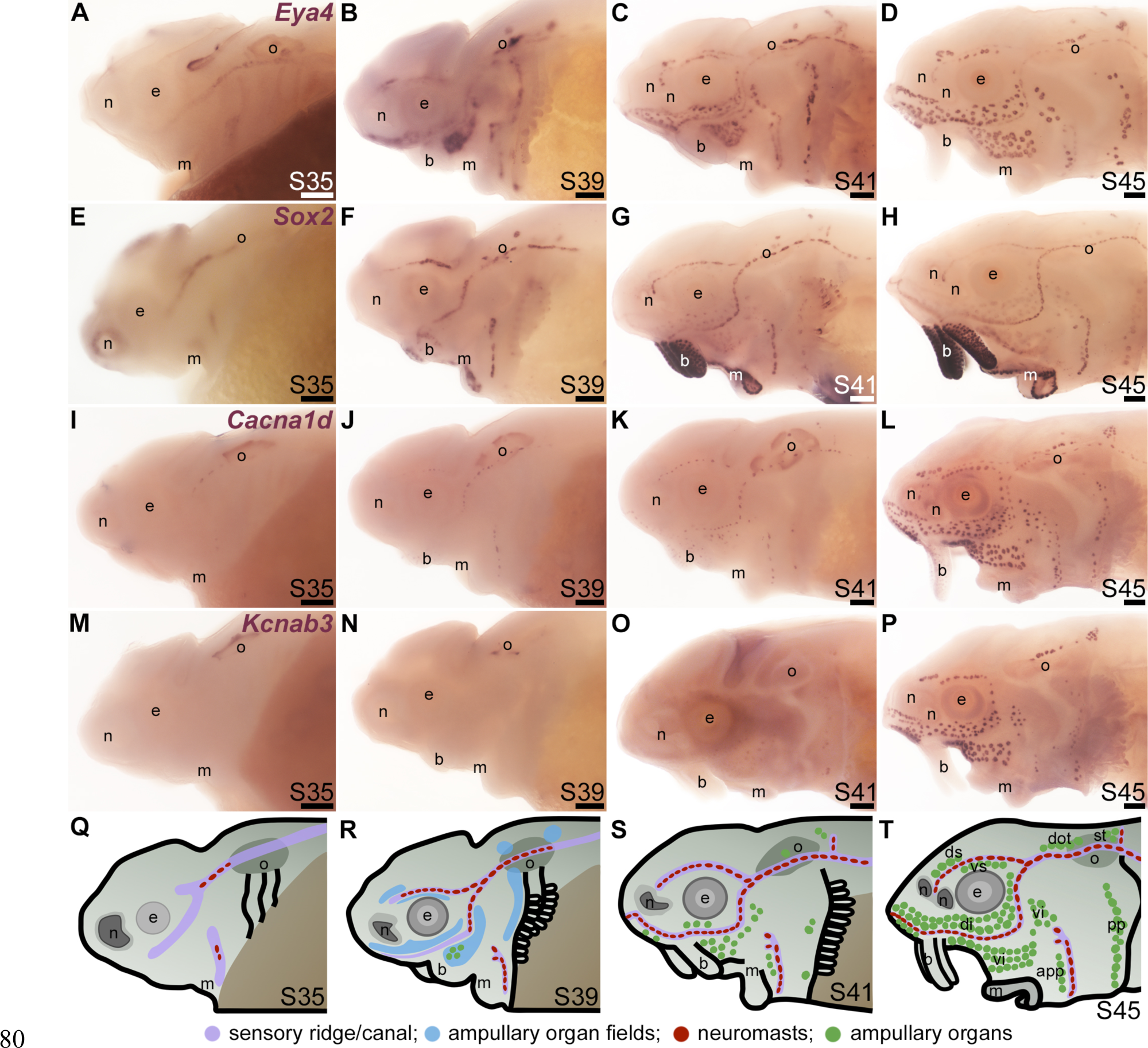
Time-course of neuromast and ampullary organ development in sterlet. *In situ* hybridization at selected stages in sterlet, from stage 35 (the stage before hatching occurs, at stage 36) to stage 45, the onset of independent feeding. (**A-D**) *Eya4* expression in sensory ridges and ampullary organ fields at stages 35 and 39 subsequently resolves into individual neuromasts and ampullary organs. (**E-H**) A paddlefish *Sox2* riboprobe reveals *Sox2* expression in sensory ridges at stage 35 that later resolves into a ring-like pattern in neuromasts, with weaker expression in ampullary organs from stage 41. Very strong expression is also seen in taste buds on the barbels and around the mouth. (**I-L**) Expression of *Cacna1d*, encoding the pore-forming alpha subunit of the voltage-gated calcium channel Cav1.3, reveals differentiated hair cells in a few neuromasts already at stage 35 in the otic line, near the otic vesicle, with increasing numbers later, and some differentiated electroreceptors already at stage 41. *Cacna1d* is also weakly expressed in taste buds, most clearly on the barbels. (**M-P**) Expression of electroreceptor-specific *Kcnab3* (encoding an accessory subunit for a voltage-gated K^+^ channel, Kvý3) shows some differentiated electroreceptors are present by stage 41, but not earlier. (**Q-T**) Schematic representation of sterlet lateral line development at stages 35, 39, 41 and 45. Abbreviations: app, anterior preopercular ampullary organ field; b, barbels; di, dorsal infraorbital ampullary organ field; dot, dorsal otic ampullary organ field; ds, dorsal supraorbital ampullary organ field; e, eye; gf, gill filaments; m, mouth; n, naris; o, otic vesicle; ppp, posterior preopercular ampullary organ field; S, stage; st, supratemporal ampullary organ field; vi, ventral infraorbital ampullary organ field; vs, ventral supraorbital ampullary organ field. Scale bar: 200 μm.

### ’Hair cell’ transcription factor genes expressed in developing ampullary organs include *Gfi1*, *Sox4* and *Sox3*

We previously showed that various transcription factor genes essential for hair cell development - *Six1*, *Eya1*, *Sox2*, *Atoh1*, *Pou4f3* (see Roccio et al., 2020; Iyer and Groves, 2021) - are expressed in developing paddlefish ampullary organs, as well as neuromasts (Modrell et al., 2011b; Modrell et al., 2011a; Butts et al., 2014; Modrell et al., 2017a). We confirm here that, in addition to *Sox2* (Figure 1E-H; Figure 2A,B), *Atoh1* and *Pou4f3* are also expressed in both types of lateral line organ in sterlet (Figure 2C-F). As noted earlier, *Sox2* was expressed more strongly in neuromasts than ampullary organs (Figure 1G,H; Figure 2A,B). However, *Atoh1* showed the converse pattern, with stronger expression in ampullary organs than in neuromasts (Figure 2C,D). These differential expression patterns were also seen at earlier stages (for *Sox2*, see Figure 1E-G; for *Atoh1*, see Supplementary Figure S2A-D).

**Figure 2.**
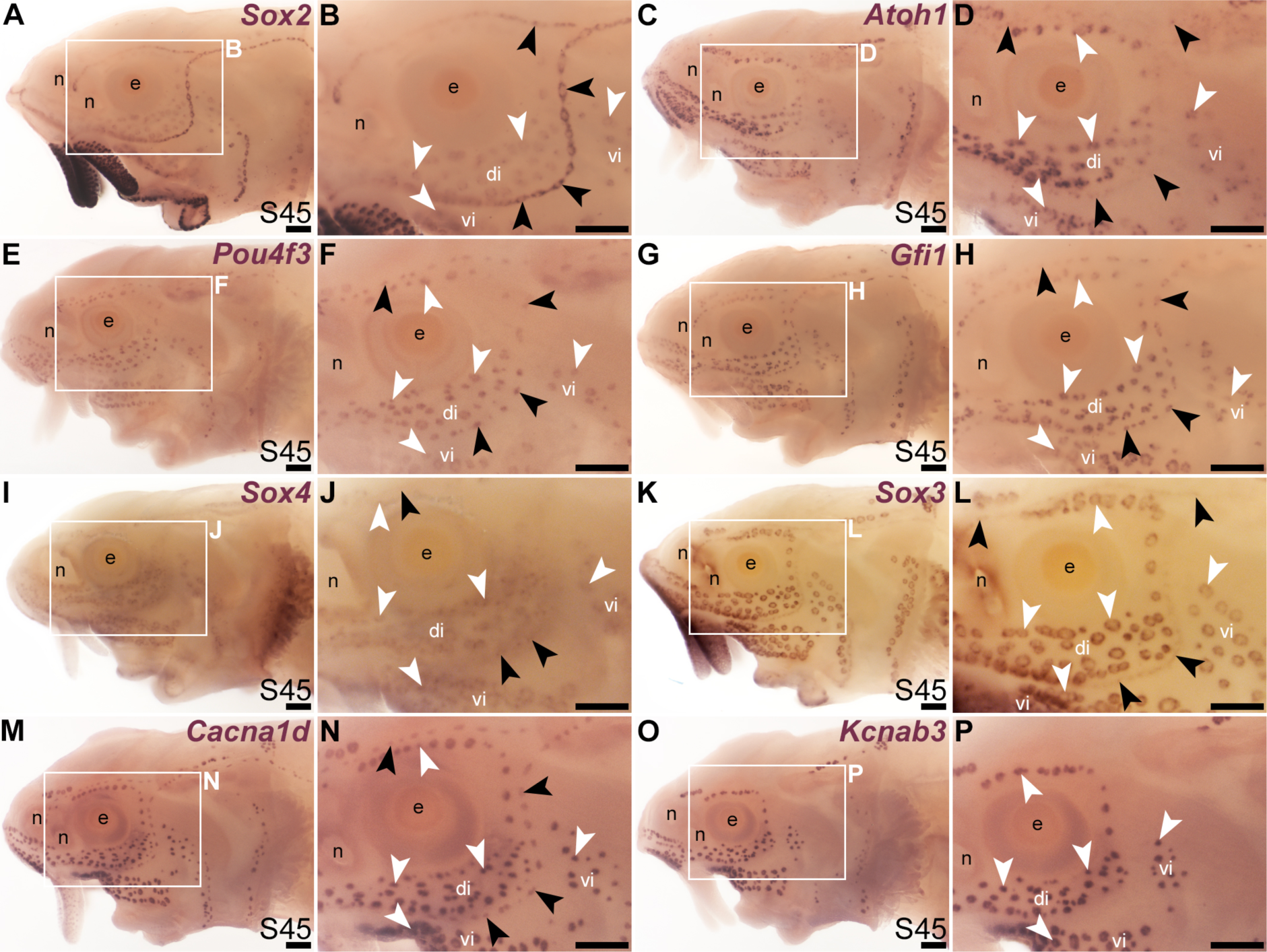
Transcription factor genes essential for hair cell development, including *Gfi1*, are expressed in ampullary organs as well as neuromasts. *In situ* hybridization in sterlet at stage 45 (the onset of independent feeding). Black arrowheads indicate examples of neuromasts; white arrowheads indicate examples of ampullary organs. (**A,B**) A paddlefish *Sox2* riboprobe reveals strong *Sox2* expression in neuromasts and weaker expression in ampullary organs (plus very strong expression in taste buds on the barbels and around the mouth). (**C,D**) *Atoh1* is expressed more strongly in ampullary organs than in neuromasts. (**E,F**) *Pou4f3* and (**G,H**) *Gfi1* are expressed in both neuromasts and ampullary organs. (**I,J**) *Sox4* is expressed in ampullary organs and very weakly in neuromasts. (**K,L**) *Sox3* expression is weaker in neuromasts than in ampullary organs (the opposite to *Sox2*; compare with A,B). (**M,N**) For comparison, the differentiated hair cell and electroreceptor marker *Cacna1d* is expressed in both neuromasts and ampullary organs (and also weakly in taste buds). (**O,P**) For comparison, the electroreceptor marker *Kcnab3* is expressed in ampullary organs only. Abbreviations: di, dorsal infraorbital ampullary organ field; e, eye; n, naris; S, stage; vi, ventral infraorbital ampullary organ field. Scale bar: 200 μm.

A key ’hair cell’ transcription factor gene whose expression we had not previously examined is the zinc-finger transcription factor gene *Gfi1* (see Roccio et al., 2020; Chen et al., 2021; Iyer and Groves, 2021; Iyer et al., 2022). *Gfi1* was 12.0-fold enriched in late-larval paddlefish operculum versus fin tissue (Modrell et al., 2017a). Sterlet *Gfi1* proved also to be expressed in developing ampullary organs, as well as neuromasts (Figure 2G,H).

In the mouse inner ear, the *SoxC* subfamily members *Sox4* and *Sox11* are co-expressed in proliferating hair cell progenitor cells and newly born hair cells, and in combination are essential for hair cell formation (Gnedeva and Hudspeth, 2015; Wang et al., 2023). Ectopic expression of either gene converts supporting cells to hair cells (Gnedeva and Hudspeth, 2015; Wang et al., 2023). A recent study showed that Sox4 confers hair-cell competence by binding lineage-specific regulatory elements and making these accessible (Wang et al., 2023). Although neither *Sox4* nor *Sox11* was present in the paddlefish lateral line-enriched gene-set (Modrell et al., 2017a), we cloned sterlet *Sox4*, which proved to be expressed in both ampullary organs and neuromasts, though more strongly in ampullary organs (Figure 2I,J). This differential expression pattern was also seen at earlier stages (Supplementary Figure S2E-H).

It has been reported that proliferative stem cells in zebrafish neuromasts express the *SoxB1* subfamily member *Sox3*, as well as *Sox2*, and that *Sox3* is important for the formation of the correct number of neuromast hair cells (preprint: Undurraga et al., 2019). A recent single-cell RNA sequencing (scRNA-seq) study showed *Sox3* expression at homeostasis in multiple neuromast support cell types (Baek et al., 2022) including central cells, the immediate precursors of regenerating hair cells (Romero-Carvajal et al., 2015; Lush et al., 2019). We had previously used a candidate gene approach for lateral line placode markers to identify *Sox3* expression in paddlefish lateral line primordia, neuromasts and also ampullary organs (Modrell et al., 2011b). (*Sox3* was 5.2-fold enriched in late-larval paddlefish operculum versus fin tissue; Modrell et al., 2017a.) As expected, sterlet *Sox3* was also expressed in both types of lateral line organ, though much more strongly in ampullary organs than in neuromasts (Figure 2K,L). Intriguingly, this was the opposite pattern to the other *SoxB1* family member, *Sox2* (Figure 2A,B). This differential expression pattern was also seen at earlier stages (Supplementary Figure S2I-L). For comparison, Figure 2M and 2N show the hair cell and electroreceptor marker *Cacna1d*, and Figure 2O and 2P show electroreceptor-specific *Kcnab3* expression.

Thus, all the key ’hair cell’ transcription factors are expressed in developing ampullary organs as well as neuromasts (although several show differing levels of expression between the two sensory organ types). These results provide further support for the hypothesis that electroreceptors evolved as transcriptionally related sister cell types to lateral line hair cells (Baker and Modrell, 2018).

### Additional transcription factor genes expressed in developing ampullary organs and neuromasts

We cloned and analysed the expression of paddlefish and/or sterlet homologues of a further 33 transcription factor genes present in the late-larval paddlefish lateral line organ-enriched dataset (Modrell et al., 2017a). (The paddlefish lateral line-enriched dataset also includes fifteen other loci assigned to specific transcription factor/co-factor genes, for which cloning and/or ISH failed in sterlet, or expression was inconsistent: *Akna*, *Barx1, Egr2, Fev*, *Fhl2*, *Fhl5*, *Litaf*, *Meis3*, *Nkx3-1*, *Not2*, *Osr1*, *Pou3f1*, *Spdef, Tbx22* and *Vgll3*.) Nine of the transcription factor genes examined were expressed in developing ampullary organs as well as neuromasts, like *Gfi1* (see previous section). One was the zinc-finger transcription factor gene *Insm2* (19.9-fold lateral line-enriched in late-larval paddlefish; Modrell et al., 2017a), which was expressed in both ampullary organs and neuromasts in paddlefish and sterlet (Figure 3A-D). However, *Insm2* expression was much stronger in ampullary organs than in neuromasts; indeed in sterlet, *Insm2* expression in neuromasts was often undetectable except in some parts of the neuromast lines (Figure 3C,D). The PRD class homeobox transcription factor gene *Otx1* (18.7-fold lateral line-enriched; Modrell et al., 2017a) similarly showed much stronger expression in ampullary organs than in neuromasts in both paddlefish and sterlet (Figure 3E-H). Also expressed in both ampullary organs and neuromasts were two other homeobox transcription factor genes, encoding TALE class Irx1 (Figure 3I,J; originally unassigned locus 111072; 8.3-fold lateral line-enriched, Modrell et al., 2017a) and LIM class Lhx6 (Figure 3K,L; originally unassigned locus 12855; 3.5-fold lateral line-enriched, Modrell et al., 2017a). The *Lhx6-like* riboprobe recognizes sequence from the 3’ untranslated region of *Lhx6-like* mRNA, as annotated in the sterlet reference genome (Du et al., 2020).

**Figure 3.**
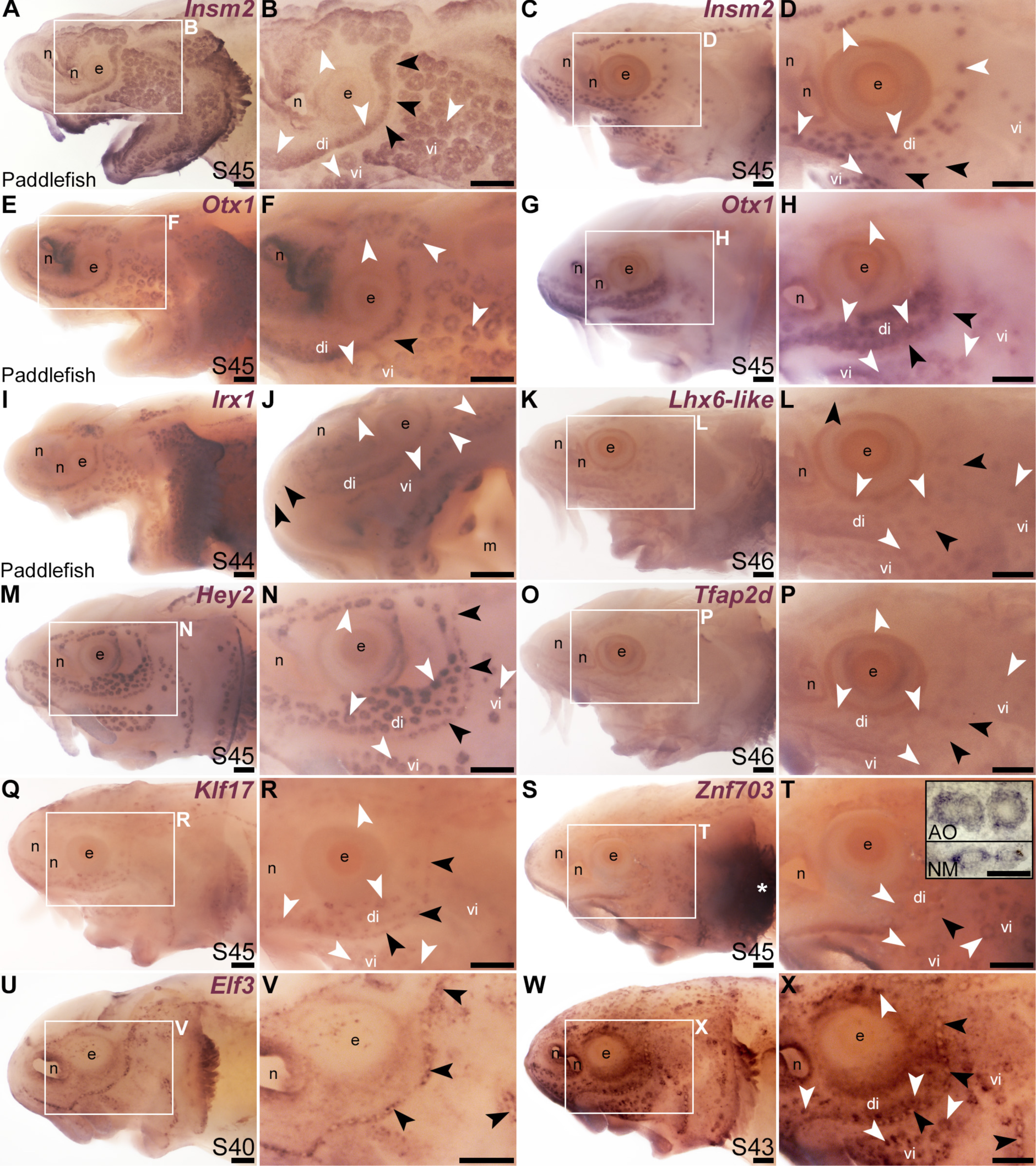
Other transcription factor genes expressed in ampullary organs and neuromasts. *In situ* hybridization in paddlefish or sterlet showing genes expressed in both ampullary organs (white arrowheads indicate examples) and neuromasts (black arrowheads indicate examples). Higher power views in each case are of the same embryo shown in the preceding panel. (**A-D**) *Insm2* at stage 45 in paddlefish (A,B) and sterlet (C,D). Neuromast expression is only detectable in some parts of the neuromast lines and is noticeably weaker than in ampullary organs. (**E-H**) *Otx1* at stage 45 in paddlefish (E,F) and sterlet (G,H). Neuromast expression is considerably weaker than ampullary organ expression (almost undetectable in paddlefish). (**I,J**) *Irx1* at stage 45 in paddlefish. Neuromast expression is weak and can be seen most clearly on the tip of the rostrum (black arrowheads in J). (**K,L**) Sterlet *Lhx6-like* at stage 46. (**M,N**) Sterlet *Hey2* at stage 45. (**O,P**) Sterlet *Tfap2* at stage 45. (**Q,R**) Sterlet *Klf17* at stage 45. (**S,T**) Sterlet *Znf703* at stage 45. Expression in lateral line organs was often hard to detect in wholemount but expression in both neuromasts (NM) and ampullary organs (AO) was clear in skinmount (examples shown in inset in T). Strong expression is seen in gill filaments (white asterisk). (**U-X**) Sterlet *Elf3*. At stage 40 (U,V), *Elf3* is expressed in a ’ring’ pattern in neuromasts, and in scattered cells in the skin. By stage 43 (W,X), *Elf3* expression is also seen in ampullary organs and more broadly throughout the skin. Abbreviations: AO, ampullary organ; di, dorsal infraorbital ampullary organ field; e, eye; m, mouth; n, naris; NM, neuromast; S, stage; vi, ventral infraorbital ampullary organ field. Scale bars: 200 μm except for inset in T: 50 μm.

The Notch target and effector gene *Hey2* (2.9-fold lateral line-enriched, Modrell et al., 2017b) was also expressed by both ampullary organs and neuromasts (Figure 3M,N). Three originally unassigned loci in the paddlefish lateral line organ-enriched dataset (Modrell et al., 2017a), all of whose closest UniProt matches had Pfam Hairy Orange and helix-loop-helix DNA-binding domains (locus 52662, 2.7-fold lateral line-enriched; locus 27975, 2.9-fold lateral line-enriched; locus 26264, 2.3-fold lateral line-enriched), proved to represent the related Notch target and effector gene *Hes5*. We had previously published the expression of *Hes5* in both ampullary organs and neuromasts in a study on the role of Notch signalling in ampullary organ versus neuromast development in paddlefish (Modrell et al., 2017b).

Expression in developing ampullary organs and neuromasts was also seen for *Tfap2d* (Figure 3O,P; 6.0-fold lateral line-enriched, Modrell et al., 2017a). This gene encodes transcription factor AP-2 delta, which is a direct activator of *Pou4f3* in retinal ganglion cell progenitors (Hesse et al., 2011; Li et al., 2016). The Krüppel-like transcription factor gene *Klf17* (2.1-fold lateral line-enriched, originally annotated in our transcriptome as *Klf4*; Modrell et al., 2017a) was also expressed in both types of lateral line organ (Figure 3Q,R), as was the zinc finger transcription factor gene *Znf703* (2.3-fold lateral line-enriched, Modrell et al., 2017a), although neuromast expression was often at the limits of detection in wholemount (Figure 3S,T). However, *Znf703* expression in neuromasts as well as ampullary organs was clear in skinmount (Figure 3T, inset). The E74-like Ets domain transcription factor gene *Elf3* (2.1-fold lateral line-enriched, Modrell et al., 2017a) showed a ’ring-like’ expression pattern in both neuromasts and ampullary organs that was clearer prior to stage 45 as general expression gradually developed throughout the skin (Figure 3U-X).

### Electrosensory-restricted cranial lateral line expression: *Irx5*, *Insm1*, *Sp5*, *Satb2*, *Mafa* and *Rorc*

Our original analysis of candidates from the late-larval paddlefish lateral line organ-enriched dataset identified the bHLH gene *Neurod4* as the first-reported transcription factor gene restricted within the paddlefish lateral line to developing ampullary organs (Modrell et al., 2017a). Here, we identified six more transcription factor genes whose cranial lateral line expression is restricted to ampullary organs. The TALE class homeobox transcription factor gene *Irx5* (1.9-fold lateral line-enriched in paddlefish, Modrell et al., 2017a) was expressed in ampullary organs but not neuromasts on the head (paddlefish: Figure 4A,B; sterlet: Figure 4C,D). However, *Irx5* expression was seen in trunk neuromasts as well as the migrating posterior lateral line primordium (Figure 4E,F). Expression of the zinc-finger transcription factor gene *Insm1* (3.1-fold lateral line-enriched, Modrell et al., 2017a) was similarly restricted to ampullary organs on the head (Figure 4G,H), but was seen in trunk neuromasts and migrating lateral line primordia (Figure 4I; compare with Sox2 immunostaining of trunk neuromasts and migrating lateral line primordia at the same stage, Figure 4J). (*Insm1* was also expressed in cells scattered throughout the skin; these are likely to be Merkel cells, which were shown to express *Insm1* in differential RNA-seq data from mouse; Hoffman et al., 2018.)

**Figure 4.**
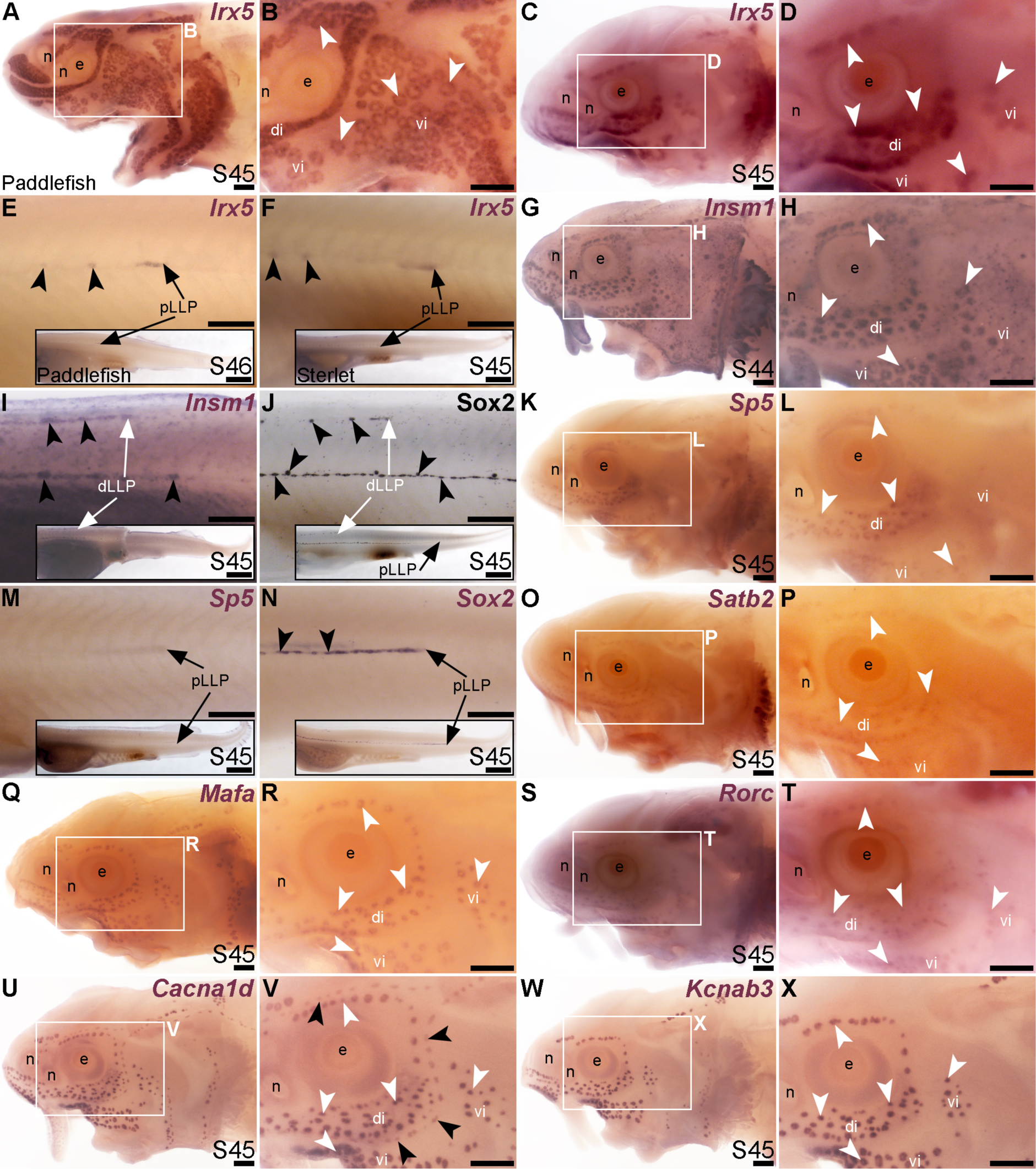
Transcription factor genes expressed in ampullary organs but not neuromasts on the head. *In situ* hybridization in paddlefish or sterlet. White arrowheads indicate examples of ampullary organs; black arrowheads indicate examples of neuromasts. (**A-F**) *Irx5* at stage 45-46 in paddlefish (A,B,E) and at stage 45 in sterlet (C,D,F; using the paddlefish riboprobe). Expression is seen in ampullary organs but not neuromasts on the head (A-D); on the trunk, expression is also visible in developing neuromasts and the migrating posterior lateral line primordium (black arrows in E,F). The insets in E,F show the position of the migrating primordia on the larval tail. (**G-I**) *Insm1* in sterlet at stages 44-45. Cranial expression is detected in ampullary organs but not neuromasts (G,H); on the trunk, expression is also seen in developing neuromasts and the migrating posterior lateral line primordia (white arrow in I shows the dorsal trunk primordium). (*Insm1* is also expressed in scattered cells throughout the skin, most likely Merkel cells.) (**J**) For comparison with panel I: Sox2 immunostaining also labels developing neuromasts and the migrating posterior lateral line primordia (white arrow: dorsal trunk line primordium; black arrow: primary posterior lateral line primordium). The inset shows the position of the migrating primordia on the larval tail. (**K-M**) *Sp5* at stage 45 in sterlet. Expression on the head is detected in ampullary organs but not neuromasts; on the trunk, weak expression is also visible in the migrating posterior lateral line primordium (arrow in K, different larva). The inset shows the position of the migrating primordium on the larval tail. (**N**) For comparison with panel M: *Sox2* expression (using a paddlefish *Sox2* riboprobe) is also seen in the migrating posterior lateral line primordium (arrow), as well as in developing neuromasts (black arrowheads). The inset shows the position of the migrating primordium on the larval tail. (**O,P**) Sterlet *Satb2* at stage 45. (**Q,R**) Sterlet *Mafa* at stage 45. (**S,T**) Sterlet *Rorc* at stage 45. (**U,V**) Sterlet *Cacna1d* at stage 45 for comparison, showing differentiated hair cells in neuromasts and electroreceptors in ampullary organs. (**W,X**) Sterlet *Kcnab3* at stage 45 for comparison, showing differentiated electroreceptors only. Abbreviations: di, dorsal infraorbital ampullary organ field; dLLP, dorsal trunk lateral line primordium; e, eye; n, naris; pLLP, posterior lateral line primordium; S, stage; vi, ventral infraorbital ampullary organ field. Scale bars: 200 μm except for insets in E,F,I,J,M,N: 1000 μm.

Another zinc-finger transcription factor gene, *Sp5* (2.6-fold lateral line-enriched, Modrell et al., 2017a), was expressed in ampullary organs but not neuromasts (Figure 4K-M), although it was expressed in the migrating posterior lateral line primordium (Figure 4M: compare with *Sox2* expression in developing neuromasts and the migrating primordium at the same stage, Figure 4N). Three other transcription factor genes were fully electrosensory-restricted: the CUT class (SATB subclass) homeobox transcription factor gene *Satb2* (4.8-fold lateral line-enriched in paddlefish, Modrell et al., 2017a), although its expression was weak (Figure 4O,P); the bZIP transcription factor gene *Mafa* (2.6-fold lateral line-enriched, Modrell et al., 2017a; Figure 4Q,R) and a retinoic acid receptor (RAR)-related orphan nuclear receptor gene, *Rorc* (14.3-fold lateral line-enriched, Modrell et al., 2017a; Figure 4S,T). For comparison with the ampullary organ-restricted cranial expression of the above-listed transcription factor genes, Figure 4U,V show *Cacna1d* expression in hair cells and electroreceptors, while Figure 4Y,X show electroreceptor-specific *Kcnab3* expression.

### *Sox8* is restricted to the mechanosensory lateral line in bony but not cartilaginous fishes

The paddlefish lateral line organ-enriched gene-set included a single *SoxE*-class high-mobility group (HMG)-box transcription factor gene, *Sox10* (3.9-fold lateral line-enriched, Modrell et al., 2017a). Comparison with the hair cell and electroreceptor marker *Cacna1d* (Figure 5A,B) and the supporting cell marker *Sox2* (Figure 5C,D) shows that paddlefish *Sox10* was not expressed within neuromasts or ampullary organs, but instead along nerves (Figure 5E,F). *Sox10* expression would be expected in nerve-associated Schwann cells, as these neural crest-derived glial cells express *Sox10* throughout their development and into the adult (see e.g., Jessen and Mirsky, 2019).

**Figure 5.**
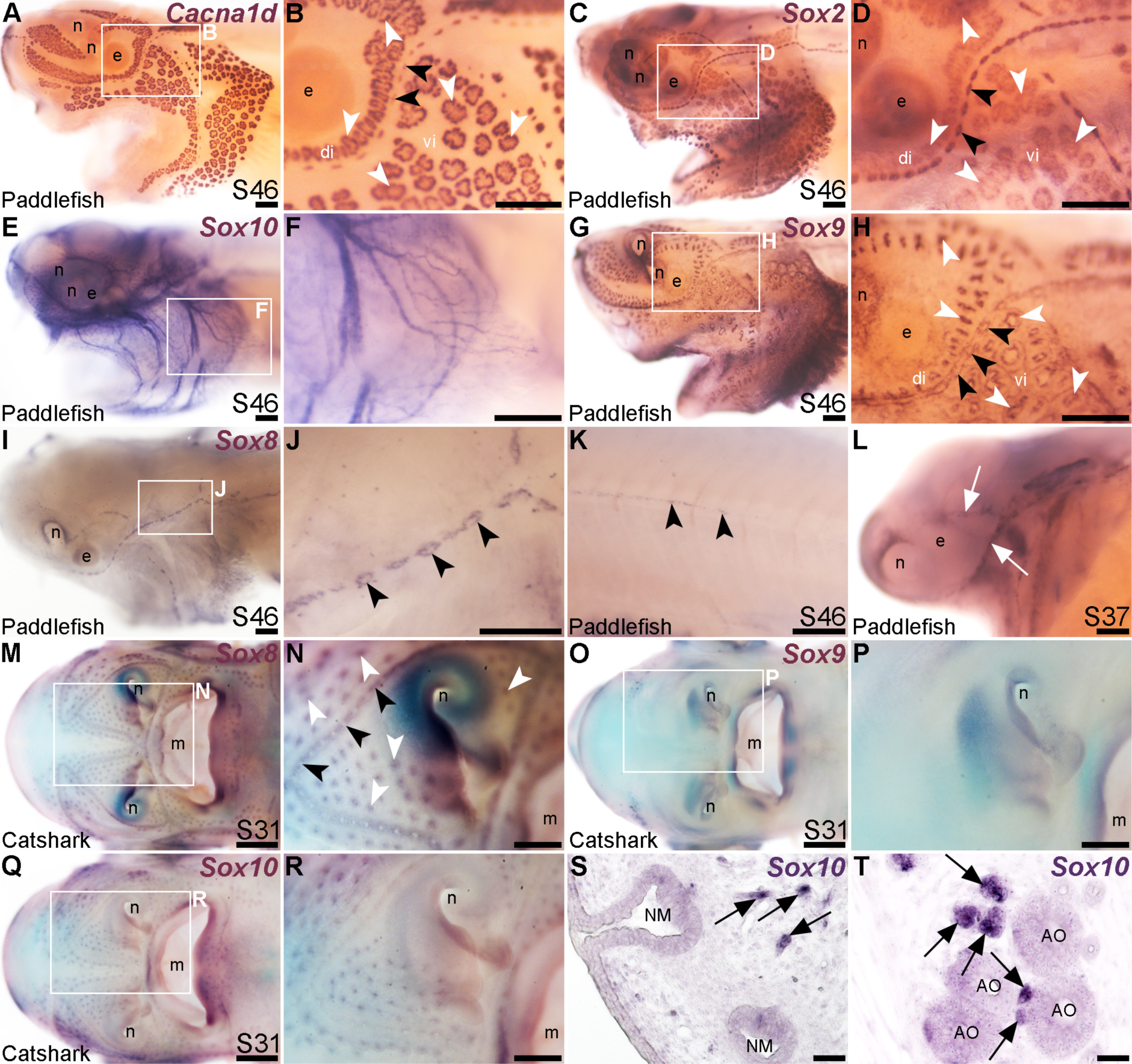
Lateral line expression of *SoxE* genes differs between chondrostean ray-finned bony fishes and cartilaginous fishes. *In situ* hybridization in late-larval paddlefish or catshark (*S. canicula*). White arrowheads indicate examples of ampullary organs; black arrowheads indicate examples of neuromasts. (**A,B**) For comparison, paddlefish *Cacna1d* expression at stage 46 reveals differentiated hair cells in neuromasts and electroreceptors in ampullary organs. (**C,D**) For comparison, paddlefish *Sox2* expression at stage 46 identifies support cells in neuromasts and ampullary organs. (**E,F**) Paddlefish *Sox10* expression at stage 46 is associated with cranial nerves, not in lateral line organs (compare with *Cacna1d* expression in panel A). (**G,H**) Paddlefish *Sox9* at stage 46 is expressed in a ’ring’-like pattern in neuromasts and ampullary organs (compare with panels A-D, especially in the ventral infraorbital ampullary organ field). (**I-K**) Paddlefish *Sox8* expression at stage 46 is seen in a ring pattern in neuromasts only (compare panel J with panels B,D), including in neuromasts developing on the trunk (K). (**L**) At stage 37, paddlefish *Sox8* is expressed in sensory ridges (white arrows). (**M,N**) Catshark *Sox8* at stage 31 is expressed in both neuromasts and ampullary organs. (**O,P**) Catshark *Sox9* expression at stage 31 is not seen in lateral line organs (compare with *Sox8* in panels O,P). (**Q-T**) Catshark *Sox10* expression at stage 31 seems to be in or near individual lateral line organs in whole-mount (Q,R). However, *in situ* hybridization on sections shows that *Sox10* is strongly expressed in cells (arrows) adjacent to neuromasts (S) and ampullary organs (T) that are likely associated with nerves. Above-background *Sox10* expression is not seen in the lateral line organs themselves. Abbreviations: di, dorsal infraorbital ampullary organ field; e, eye; m, mouth; n, naris; S, stage; vi, ventral infraorbital ampullary organ field. Scale bars: A-L,N,P,R, 200 μm; M,O,Q, 500 μm; S,T, 10 μm.

Another *SoxE*-class HMG-box transcription factor gene, *Sox8*, was previously reported to be expressed in developing ampullary organs in a shark, *Scyliorhinus canicula* (Freitas et al., 2006). Given this, we also cloned *Sox8* and the remaining *SoxE* class gene, *Sox9*, to test their expression in paddlefish. *Sox9* was expressed at late-larval stages in both neuromasts and ampullary organs with a ’ring-like’ distribution (Figure 5G,H). This is also consistent with Sox9 expression in the developing mouse inner ear, where initial broad expression becomes restricted to supporting cells, co-expressed with Sox2 (Mak et al., 2009; Jan et al., 2021).

Paddlefish *Sox8* expression, in contrast to *Sox9*, was restricted to neuromasts at late-larval stages (Figure 5I,J), also in a ring pattern suggestive of supporting cells rather than centrally clustered hair cells (compare neuromast expression of *Sox8* in Figure 5J with *Cacna1d* in hair cells in Figure 5B and *Sox2* expression in supporting cells in Figure 5D). *Sox8* was also expressed in neuromasts on the trunk (Figure 5K), and at earlier stages, in the central region of sensory ridges where neuromasts form (Figure 5L).

The mechanosensory lateral line-restricted expression of paddlefish *Sox8* contrasts with the reported expression of *Sox8* in shark ampullary organs (Freitas et al., 2006). To test this further, we cloned all three *SoxE* genes from the lesser-spotted catshark (*Scyliorhinus canicula*). We confirmed that *Sox8* is expressed in shark ampullary organs, as previously reported (Freitas et al., 2006), as well as in neuromasts (Figure 5M,N), unlike the neuromast-specific *Sox8* expression seen in the late-larval paddlefish (Figure 5K,L). *Sox9* was not expressed in shark lateral line organs at all (Figure 5O,P), in striking contrast to paddlefish *Sox9* expression in both neuromasts and ampullary organs (Figure 5G,H). The only conserved *SoxE* lateral line expression pattern between catshark and paddlefish was that of *Sox10*, which ISH on sections confirmed to be restricted to axon-associated Schwann cells (Figure 5Q-T). Overall, these data reveal lineage-specific differences in *SoxE* transcription factor gene expression within late-larval lateral line organs in a ray-finned chondrostean fish versus a cartilaginous fish.

### *Foxg1* is restricted to the mechanosensory lateral line

The winged-helix transcription factor gene *Foxg1* was 11.4-fold enriched in late-larval (stage 46) paddlefish operculum vs. fin (Modrell et al., 2017a). *Foxg1* proved to be restricted to the mechanosensory lateral line in ray-finned chondrostean fishes. For comparison, Figure 6A-D show Sox2 protein expression in supporting cells in neuromasts and (more weakly) in ampullary organs in late-larval paddlefish (Figure 6A,B) and sterlet (Figure 6C,D). Sox2 immunostaining also labels taste buds, and individual cells scattered throughout the skin (most likely Merkel cells, which express Sox2 in zebrafish, as well as mouse; Brown et al., 2023; Bardot et al., 2013; Lesko et al., 2013; Perdigoto et al., 2014). Paddlefish *Foxg1* expression in the lateral line system at stage 45 (Figure 6E,F) was restricted to neuromast lines, but excluded from the central domain of individual neuromasts where hair cells are found (compare Figure 6F with Sox2 in Figure 6B). (*Foxg1* expression was also seen in the olfactory system, as expected; e.g., Kawauchi et al., 2009.) Paddlefish *Foxg1* was expressed in the migrating posterior lateral line primordium on the trunk, and in trunk neuromasts deposited by the primordium (Figure 6G; compare with Sox2 immunostaining at the same stage, Figure 6H). Sterlet *Foxg1* was expressed in the same pattern at stage 45 as in paddlefish (Figure 6I-K; Figure 6L shows Sox2 immunostaining in the sterlet posterior lateral line primordium and trunk neuromasts for comparison with sterlet *Foxg1* expression in Figure 6K). Analysis at earlier stages in sterlet showed that *Foxg1* expression was restricted to the central zone of sensory ridges where neuromasts form (Figure 6M-R). Thus, *Foxg1* expression in the developing lateral line system is restricted to the mechanosensory division, although it seems to be excluded from differentiated hair cells.

**Figure 6.**
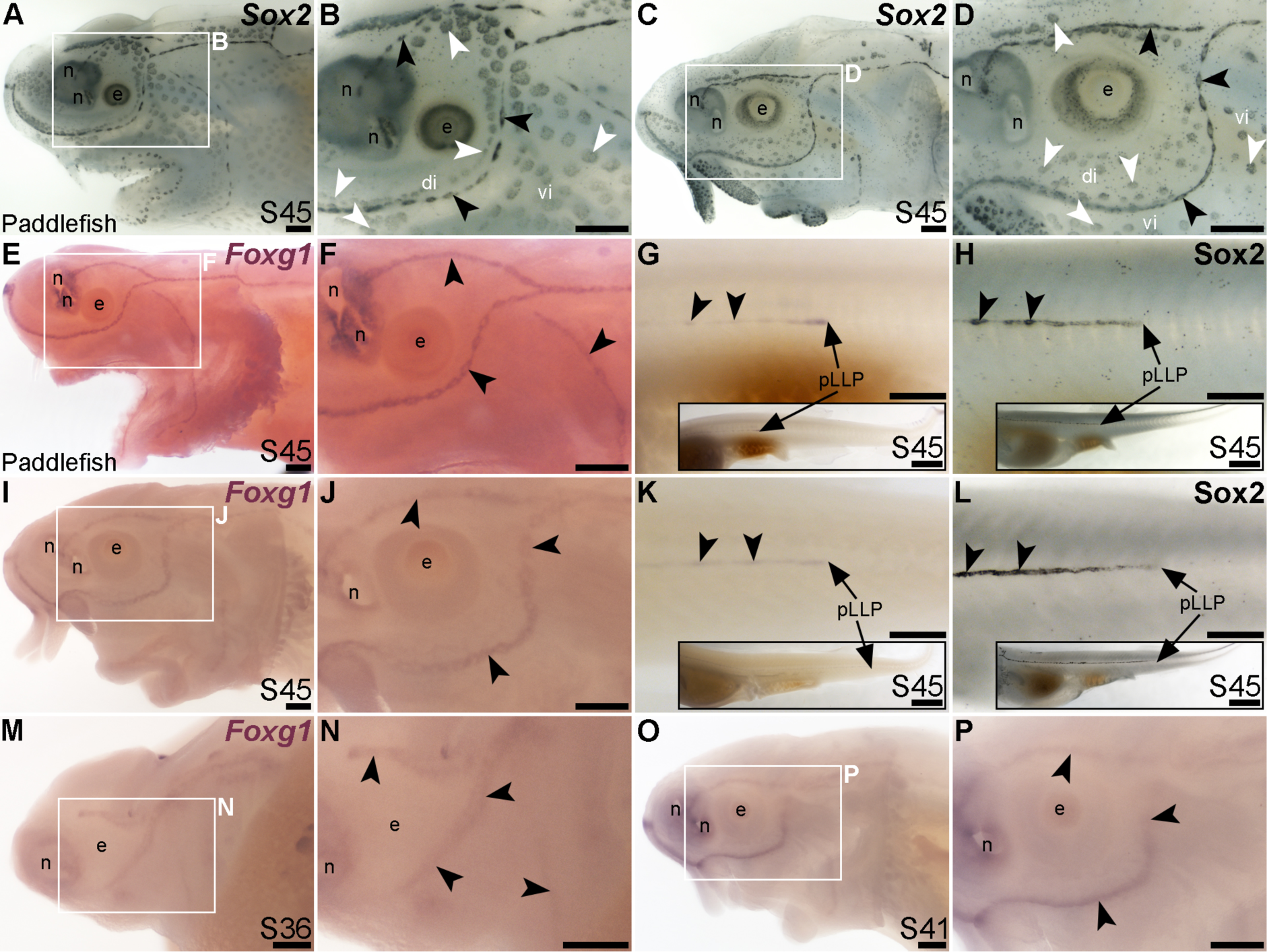
*Foxg1* is mechanosensory-restricted within the developing lateral line system. Black arrowheads indicate examples of neuromasts; white arrowheads indicate examples of ampullary organs. (**A-D**) For comparison, Sox2 immunostaining at stage 45 in paddlefish (A,B) and sterlet (C,D) shows support cells in neuromasts (stronger staining) and ampullary organs (weaker staining). (Strong Sox2 expression is also seen in taste buds on the barbels and around the mouth, and in scattered cells in the skin, most likely Merkel cells.) (**E-G**) *In situ* hybridization for *Foxg1* at stage 45 in paddlefish, showing a ring-like expression pattern in the neuromast lines (compare panel F with paddlefish Sox2 in B), as well as expression in the migrating posterior lateral line primordium (arrow in G, different larva) and developing trunk neuromasts (black arrowheads). Expression is also seen in the nares. The inset in G shows the position of the migrating primordium on the larval tail. (**H**) For comparison with G, Sox2 immunostaining on the trunk at stage 45 in paddlefish shows the migrating posterior lateral line primordium (arrow) and developing neuromasts. The inset shows the position of the migrating primordium on the larval tail. (**I-K**) *In situ* hybridization for *Foxg1* at stage 45 in sterlet similarly shows a ring pattern in neuromasts (compare panel J with sterlet Sox2 in D), as well as expression in the migrating posterior lateral line primordium (arrow in K, different larva) and developing trunk neuromasts (black arrowheads). Expression is also seen in the nares. The inset in K shows the position of the migrating primordium on the larval tail. (**L**) For comparison with K, Sox2 immunostaining on the trunk at stage 45 in sterlet shows weak expression in the migrating posterior lateral line primordium (arrow) and developing neuromasts. The inset shows the position of the migrating primordium on the larval tail. (**M-P**) *In situ* hybridization for sterlet *Foxg1* at stage 36 (M,N) and stage 42 (O,P) shows ring-like expression already in developing neuromasts in sensory ridges, and no expression in developing ampullary organ fields (compare with sterlet *Eya4* expression at stages 35 and 41 in Figure 1A,C). Abbreviations: di, dorsal infraorbital ampullary organ field; e, eye; n, naris; pLLP, posterior lateral line primordium; S, stage; vi, ventral infraorbital ampullary organ field. Scale bars 200 μm except for insets in G,H,K,L: 1000 μm.

### Mechanosensory-restricted lateral line expression of *Hmx2*, *Isl1* and *Rorb*

The NKL class homeobox transcription factor gene *Hmx2* (also known as *Nkx5-2*), which was 5.8-fold lateral line-enriched in late-larval paddlefish (Modrell et al., 2017a), also proved to be restricted to the mechanosensory lateral line. For comparison, Figure 7A,B show Sox2 immunostaining at stage 45. At this stage, *Hmx2* was expressed in a ring-like pattern at the outer edge of developing neuromasts (Figure 7C,D; compare with Sox2 in Figure 7A,B). However, no expression was seen in ampullary organs (Figure 7C,D), confirmed in skinmounts after post-ISH Sox2 immunostaining to identify ampullary organs (Figure 7E). As an aside, *Hmx2* expression was also seen in scattered skin cells (Figure 7C-E). *Hmx2* was not among the genes reported in a differential RNA sequencing study of adult mouse Merkel cells (Hoffman et al., 2018). However, examination of sterlet skinmounts after post-ISH Sox2 immunostaining suggested the *Hmx2*-expressing skin cells may co-express Sox2 (Figure 7E), which would support their being Merkel cells. Alternatively, other scRNA-seq studies in mouse and human have shown that *Hmx2* is expressed by tuft (brush) cells in gut and airway epithelia (Haber et al., 2017; Deprez et al., 2020), so it is possible the *Hmx2*-expressing skin cells in paddlefish are chemosensory epithelial cells, like tuft cells (Kotas et al., 2023).

**Figure 7.**
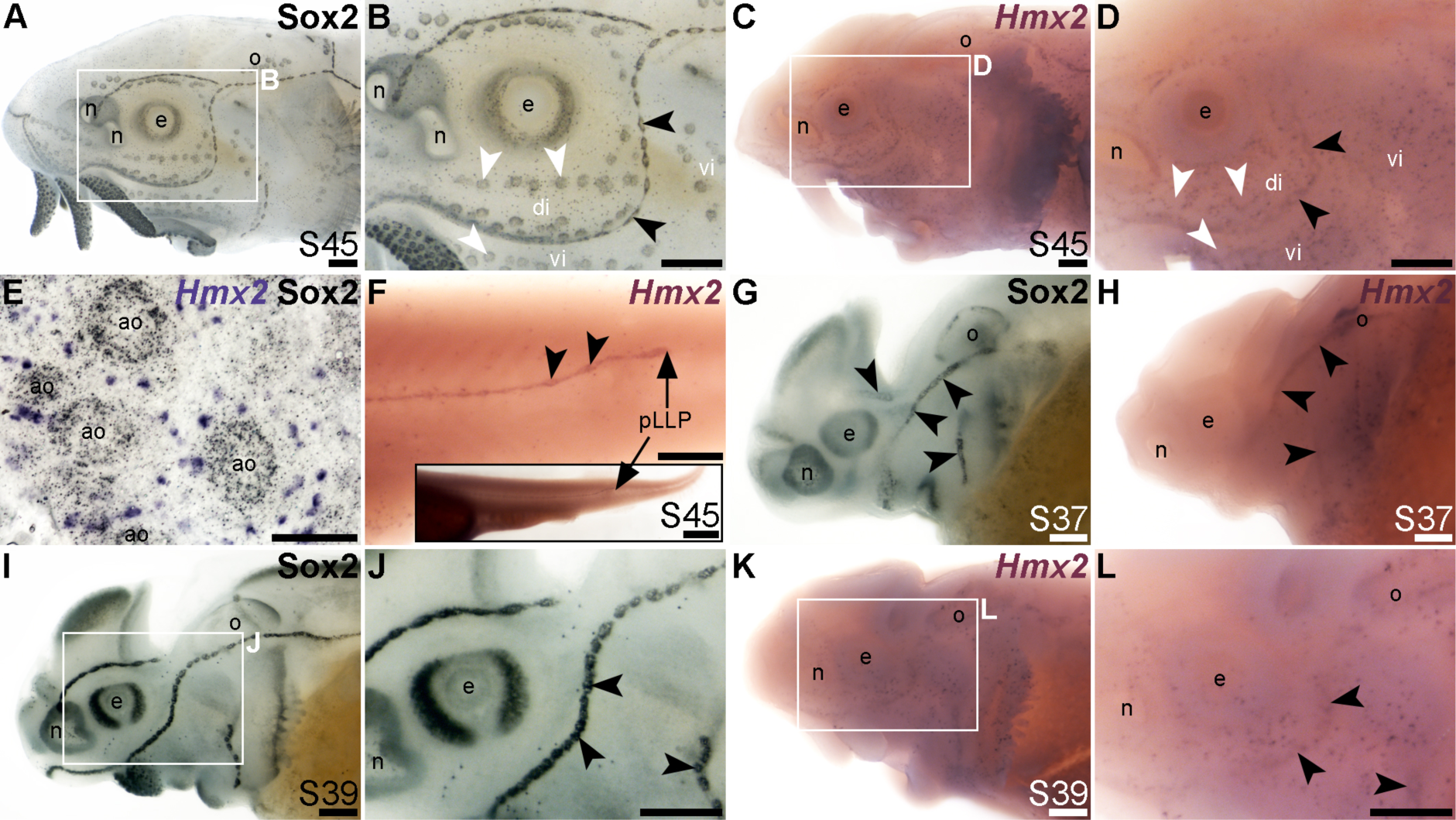
*Hmx2* is mechanosensory-restricted within the developing lateral line system. Black arrowheads indicate examples of neuromasts; white arrowheads indicate examples of ampullary organs. (**A,B**) For comparison, Sox2 immunostaining in sterlet is shown at stage 45. Sox2 labels developing neuromasts (stronger staining) and scattered cells in the skin, most likely Merkel cells, as well as taste buds on the barbels and around the mouth. Expression is also seen in ampullary organs (weaker than in neuromasts). A patch of Sox2 expression at the spiracular opening (first gill cleft) may represent the spiracular organ. (**C-F**) *In situ* hybridization for sterlet *Hmx2* at stage 45 in wholemount (C,D), *Hmx2* is weakly expressed in a ring-like pattern in the neuromast lines, as well as in scattered cells in the skin, but appears to be absent from ampullary organs (compare D with Sox2 expression in B). A skinmount (E) with several ampullary organs from a stage 45 embryo revealed by post-ISH immunostaining for Sox2 (black metallographic deposits) confirms that ampullary organs do not express *Hmx2* (purple). (The scattered *Hmx2*-positive skin cells may co-express Sox2, suggesting they are likely to be Merkel cells.) *Hmx2* is also expressed in the trunk neuromast line (F), including the migrating posterior lateral line primordium (black arrow in J). The inset shows the position of the migrating primordium on the larval tail. (**G**) For comparison with H, Sox2 immunostaining at stage 37 labels developing neuromasts. Expression is also seen in the nasal capsule and otic vesicle, as well as eye. (**H**) *Hmx2* expression at stage 37 is detected in neuromast lines, as well as in the otic vesicle. (**I,J**) For comparison with K and L, Sox2 immunostaining at stage 39 labels developing neuromasts and scattered cells in the skin, most likely Merkel cells, as well as taste buds on the barbels. Expression is also seen in the nasal capsule and the eye. (**K,L**) *Hmx2* expression at stage 39 is detected in neuromast lines and scattered cells in the skin. Abbreviations: ao, ampullary organ; di, dorsal infraorbital ampullary organ field; e, eye; n, naris; o, otic vesicle; pLLP, posterior lateral line primordium; S, stage; vi, ventral infraorbital ampullary organ field. Scale bars: 200 μm except for panel E: 50 μm and inset in F, 1000 μm.

*Hmx2* was also expressed in the migrating posterior lateral line primordium at stage 45 (Figure 7F), like *Foxg1* (Figure 6K) and Sox2 (Figure 6L). Analysis at earlier stages, with Sox2 immunostaining for comparison (Figure 7G-L), showed that *Hmx2* was weakly expressed in neuromast lines (and in the otic vesicle) at stage 37 (Figure 7H; compare with Sox2 in Figure 7G) and at stage 39 (Figure 7K,L; compare with Sox2 in Figure 7I,J).

The LIM class homeobox transcription factor Isl1 was recently reported to promote a more complete conversion by Atoh1 of mouse cochlear supporting cells to hair cells than does Atoh1 alone (Yamashita et al., 2018). The only LIM homeobox genes in the paddlefish lateral line organ-enriched dataset (Modrell et al., 2017a) are *Lhx3*, which we previously reported to be expressed in ampullary organs as well as neuromasts (Modrell et al., 2017a); *Lhx6-like* (originally unassigned locus 12855), with the same expression pattern (Figure 3K), and *Lhx8*, which proved to be expressed in gill filaments, not lateral line organs (Supplementary Figure S3J). Nevertheless, given the demonstrated role for Isl1 in promoting cochlear hair cell formation (Yamashita et al., 2018), we cloned sterlet *Isl1*. For comparison, Figure 8A,B show Sox2 immunostaining at stage 45, and Figure 8C,D show *Cacna1d*-expressing hair cells and electroreceptors at stage 45. At this stage, *Isl1* expression was weak, but restricted within the lateral line system to neuromasts (Figure 8E,F). Examination at earlier stages revealed no detectable expression at stage 36 (Figure 8G,H) and neuromast-restricted expression at stages 39 and 40, when *Isl1* seemed to be more strongly expressed than at stage 45 (Figure 8I-L).

**Figure 8.**
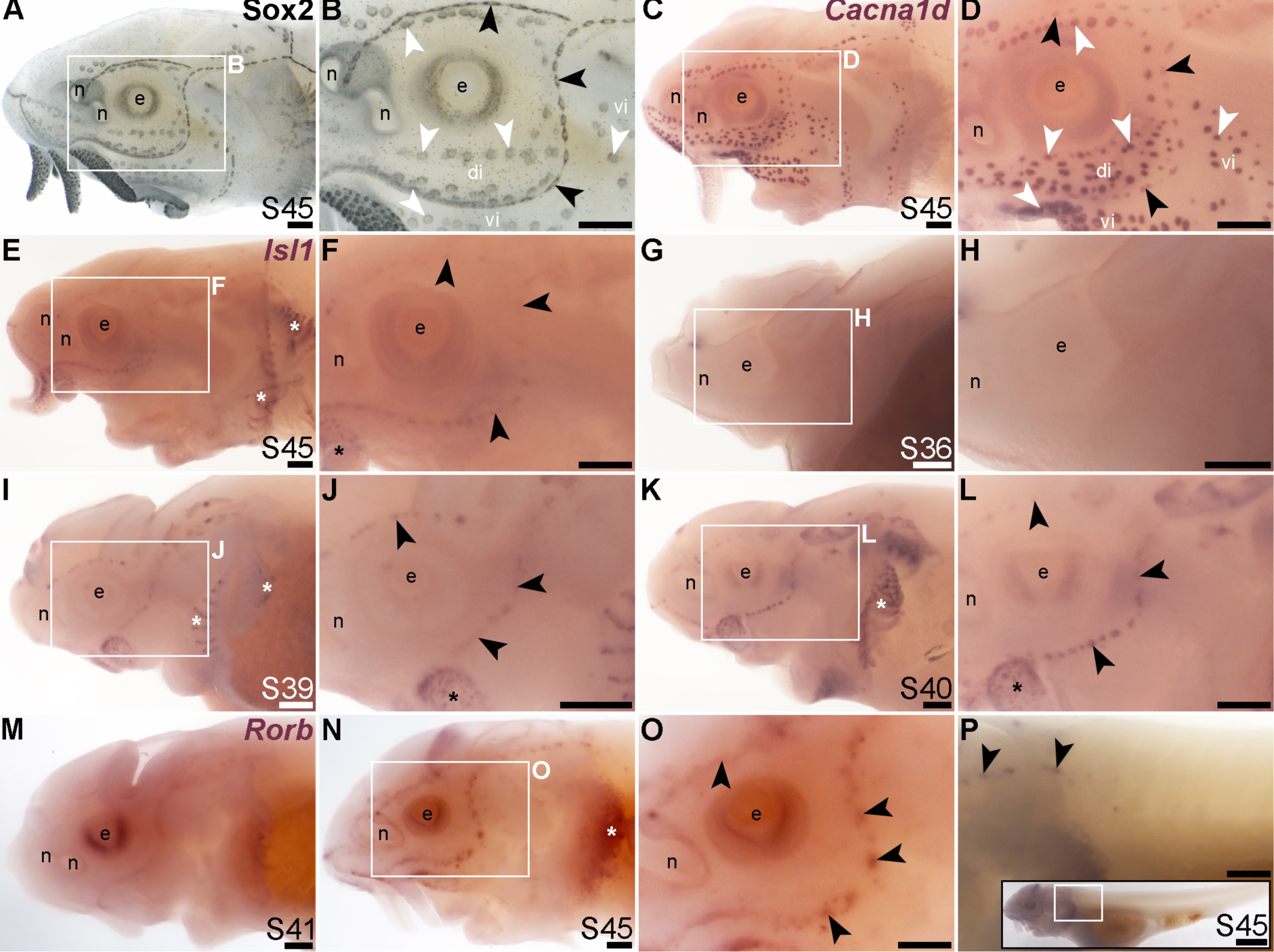
*Isl1* and *Rorb* are mechanosensory-restricted within the developing lateral line system. Black arrowheads indicate examples of neuromasts; white arrowheads indicate examples of ampullary organs. (**A,B**) For comparison, Sox2 immunostaining in sterlet is shown at stage 45. Sox2 labels developing neuromasts (stronger staining) and scattered cells in the skin, most likely Merkel cells, as well as taste buds on the barbels and around the mouth. Expression is also seen in ampullary organs (weaker than in neuromasts). A patch of Sox2 expression at the spiracular opening (first gill cleft) may represent the spiracular organ. (**C,D**) For comparison, *in situ* hybridization for sterlet *Cacna1d* at stage 45 shows expression in hair cells in neuromasts and electroreceptors in ampullary organs (and weak expression in taste buds on the barbels). (**E,F**) *In situ* hybridization for sterlet *Isl1* at stage 45 shows weak spots of expression in neuromasts but not ampullary organs (compare with *Cacna1d* in C,D). Stronger expression is seen in taste buds on the barbels (black asterisk) and in gill filaments (white asterisk). (**G-L)** *In situ* hybridization for sterlet *Isl1* at earlier stages shows no expression at stage 36 (G,H), and expression in neuromasts (but not ampullary organs) at stage 39 (I,J), and stage 40 (K,L). From stage 39, *Isl1* expression is also seen in taste buds on the barbels (asterisk in J) and in gill filaments. (**M-P**) *In situ* hybridization for sterlet *Rorb* at stage 41 (M), and stage 45 (N-P) shows cranial neuromast-specific expression within the lateral line (i.e., without expression in either trunk neuromasts or the migrating posterior lateral line primordium). Abbreviations: di, dorsal infraorbital ampullary organ field; e, eye; n, naris; pLLP, posterior lateral line primordium; S, stage; vi, ventral infraorbital ampullary organ field. Scale bars: 200 μm.

Finally, we identified the RAR-related orphan nuclear receptor beta gene, *Rorb* (6.2-fold lateral line-enriched; Modrell et al., 2017a), as being restricted to cranial neuromasts, with no detectable expression in ampullary organs or trunk neuromasts (Figure 8M-P). The onset of *Rorb* expression in neuromasts was later even than *Isl1*, starting only at stage 41 (Figure 8M,N). It was intriguing to see the mutually exclusive expression of *Rorb* in cranial neuromasts (Figure 8O,P) and *Rorc* in ampullary organs (Figure 4O,P).

Taken together, we have identified *Sox8*, *Foxg1*, *Hmx2*, *Isl1* and *Rorb* as the first-reported transcription factor genes restricted to the mechanosensory division of the lateral line system in ray-finned fishes. *Sox8* and *Foxg1* are expressed in the central zone of sensory ridges where neuromasts form and maintained in neuromasts, though apparently excluded from differentiated hair cells. *Hmx2* is expressed in sensory ridges and retained in neuromasts, whereas *Isl1* and *Rorb* are restricted to neuromasts (specifically cranial neuromasts, for *Rorb*) as early as they can be detected.

The remaining transcription factor genes from the paddlefish lateral line organ-enriched gene-set that we examined proved not to be expressed in lateral line organs, but instead, e.g., in ectoderm around ampullary organs (Supplementary Figure S3A-D: *Ehf*, *Foxi2* and *Nkx2-3*), or at the edge of the operculum, in taste buds and/or in developing gill filaments (Supplementary Figure S3E-P: *Foxe1*, *Foxl2*, *Gcm2*, *Hoxa2*, *Lhx8*, *Pou3f4*, *Sim2*, *Tbx1*, *Tlx1*, *Tlx2* and *Rax2*).

## Discussion

In this study, we used our paddlefish lateral line organ-enriched gene-set (generated from differential bulk RNA-seq at late larval stages; Modrell et al., 2017a), together with a candidate gene approach, to identify 23 novel transcription factor genes expressed in developing lateral line organs in sterlet and/or paddlefish. These data, together with our previous work in paddlefish (Modrell et al., 2011b; Modrell et al., 2011a; Modrell et al., 2017b; Modrell et al., 2017a), suggest extensive conservation of molecular mechanisms involved in electrosensory and mechanosensory lateral line organ development. However, they also reveal a set of transcription factor genes with restricted expression that may be involved in the development of mechanosensory versus electrosensory organs. Of the 40 transcription factor genes with validated expression during lateral line organ development in paddlefish and/or sterlet, 28 (70%) were expressed in both ampullary organs and neuromasts (Table 1). These include the key ’hair cell’ transcription factor genes *Six1*, *Eya1*, *Sox2*, *Atoh1*, *Pou4f3* and *Gfi1* (see Roccio et al., 2020; Chen et al., 2021; Iyer and Groves, 2021; Iyer et al., 2022). We also identified seven electrosensory-restricted and five mechanosensory-restricted transcription factor genes (Table 1), as discussed further below.

**Table 1.**
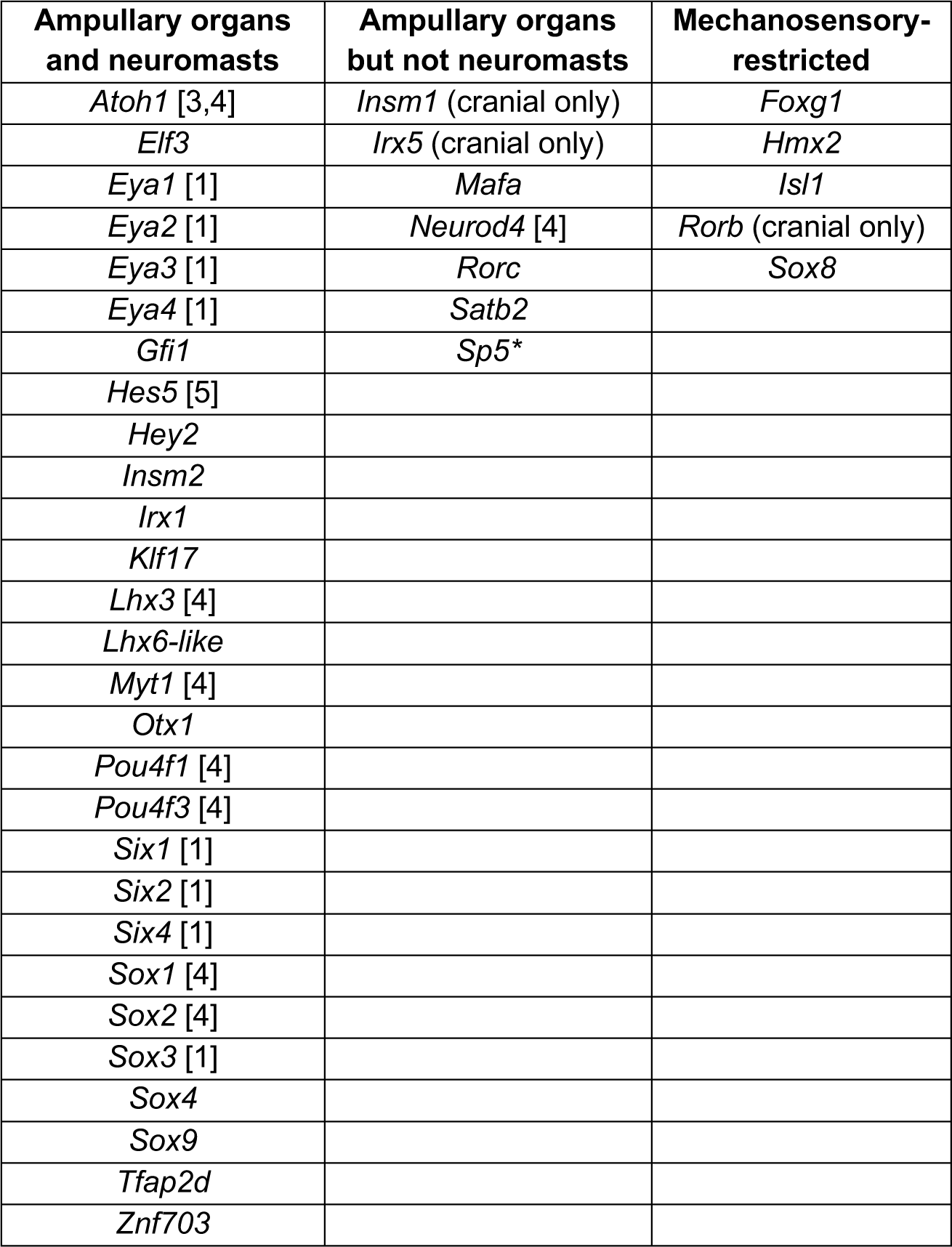
Transcription factor genes expressed in developing lateral line organs in paddlefish and/or sterlet. Lateral line expression was reported either in this study or in previous papers, denoted by numbers in brackets: [1] Modrell et al. (2011b); [2], Modrell et al. (2011a); [3] Butts et al. (2014); [4] Modrell et al. (2017a); [5] Modrell et al. (2017b).

While this work was ongoing, a differential RNA-seq study of regenerating ampullary organs and neuromasts in the Siberian sturgeon (*Acipenser baerii*) was published (Wang et al., 2020). This study compared dissected tissue samples containing stage 45 ampullary organs or neuromasts relative to general epidermis, which identified 2074 lateral line organ-enriched genes, of which 1418 were shared by ampullary organs and neuromasts; 539 were ampullary organ-enriched, and 117 were neuromast-enriched (Wang et al., 2020). The ’common’ stage 45 lateral line-organ dataset from the Siberian sturgeon (Wang et al., 2020) included many of the candidate genes encoding transcription factors and differentiation markers whose expression we have validated in both ampullary organs and neuromasts at stage 45-46 in paddlefish and/or sterlet, e.g. *Six1*, *Eya1*, *Atoh1*, *Pou4f3*, *Gfi1*, *Otof,* and *Cacna1d* (Modrell et al., 2011b; Modrell et al., 2011a; Modrell et al., 2017a; this study). The ampullary organ-enriched dataset from the Siberian sturgeon (Wang et al., 2020) included *Sp5*, which we identified here in sterlet as ampullary organ-restricted, but also *Hmx2* and *Rorb*, which we identified here as neuromast-restricted. Conversely, the neuromast-enriched dataset (Wang et al., 2020) included *Insm1*, which we found in sterlet to be ampullary organ-restricted on the head (though expressed in trunk neuromasts). Furthermore, the ’common’ dataset (Wang et al., 2020) included genes whose expression in sterlet and paddlefish was either ampullary organ-specific (e.g., the voltage-gated K^+^ channel genes *Kcna5* and *Kcnab3*, as well as *Neurod4*; this study; Modrell et al., 2017a), or mechanosensory-specific (e.g., *Foxg1* and *Isl1*; this study).

Overall, we think that the stage 45 Siberian sturgeon tissue dissections (Wang et al., 2020) were unable to separate ampullary organs and neuromasts completely. Like our own stage 46 paddlefish lateral line organ-enriched dataset (Modrell et al., 2017a), the stage 45 Siberian sturgeon datasets are not exhaustive (Wang et al., 2020): some of the genes whose expression we have validated in stage 45-46 sterlet and/or paddlefish lateral line organs were missing (e.g., *Satb2*, *Sox2* and *Sox8*; this study; Modrell et al., 2017a). Nevertheless, this differential RNA-seq study in late-larval Siberian sturgeon embryos (Wang et al., 2020) provides an invaluable, independent resource from which to identify additional candidate genes for future validation and functional investigation *in vivo*.

### Conserved molecular mechanisms are likely involved in the formation of electroreceptors and hair cells

In this study, we identified 12 novel transcription factor genes expressed in both types of lateral line organ in chondrostean ray-finned fishes, consistent with conservation of molecular mechanisms. In particular, we highlight *Gfi1* (see Roccio et al., 2020; Chen et al., 2021; Iyer and Groves, 2021; Iyer et al., 2022) as being another key ’hair cell’ transcription factor gene expressed in developing ampullary organs as well as neuromasts, together with *Atoh1*, *Pou4f3* and *Six1* (Modrell et al., 2011a; Modrell et al., 2017a). *Gfi1*-deficient hair cells fail to mature and also upregulate neuronal differentiation genes such as *Neurod1* and *Pouf41* (and *Insm1*, which is important for otic neurogenesis as well as outer hair cell formation; Lorenzen et al., 2015), suggesting that a key function of Gfi1 in hair cells is to repress neuronal genes that are initially also expressed in hair cell progenitors (Matern et al., 2020). Gfi1 also acts indirectly to increase Atoh1 transcriptional activity by forming part of a transcriptional complex with Atoh1 and E proteins in which neither Atoh1 nor Gfi1 binds the other directly and Gfi1 does not bind DNA (Jen et al., 2022). Given the shared expression in ampullary organs and neuromasts, it seems likely that Gfi1 plays these roles in both developing electroreceptors and hair cells. Intriguingly, however, *Insm2* was recently reported as a direct target of both Atoh1 and Gfi1 in mouse cochlear hair cells, and one of only a handful of genes (including *Atoh1* itself) to be repressed by Gfi1 during hair-cell maturation (Jen et al., 2022). Repression of *Insm2* by Gfi1 in mature hair cells, but not electroreceptors, could explain the much weaker expression of *Insm2* that we saw in neuromasts versus ampullary organs at stage 45 (the onset of independent feeding). This suggests the existence of both shared and divergent functions of the same transcription factor within hair cells versus electroreceptors.

### Six novel transcription factor genes with ampullary organ-restricted cranial expression

We have identified six novel transcription factor genes expressed in developing ampullary organs but not cranial neuromasts in chondrostean ray-finned fishes, in addition to previously published *Neurod4* (Modrell et al., 2017a). Two of these, *Irx5* and *Satb2*, encode homeodomain transcription factors. In *C. elegans*, unique combinations of homeodomain transcription factors define all 118 neuron classes (Reilly et al., 2020) (also see Vidal et al., 2022). Hence, members of this class of transcription factors are potentially good candidates to be involved in controlling divergent fate specification and/or maintenance in closely related cell types.

*Irx5* is required for the terminal differentiation of a subset of cone bipolar cells in the mouse retina (Cheng et al., 2005). In mouse and chicken, *Irx5* is expressed within the otic vesicle epithelium, including some prospective sensory patches (Bosse et al., 2000; Cardeña-Núñez et al., 2016). However, by stage 34 in chicken (embryonic day 8), when hair cells are fully differentiated, *Irx5* is not expressed in any sensory patch, unlike some other *Irx* family members (Cardeña-Núñez et al., 2016). In mouse, chicken and zebrafish, *Irx5* (together with other *Irx* family members) is expressed in otic placode-derived neurons (Bosse et al., 2000; Houweling et al., 2001; Cardeña-Núñez et al., 2016; Lecaudey et al., 2005). In zebrafish, the only reported expression of *Irx5a* or *Irx5b* in the lateral line system is that of *Irxa* in the secondary posterior lateral line primordium (prim II) (Lecaudey et al., 2005). This migrates later than the primary posterior lateral line primordium and contributes post-embryonically to lateral and dorsal branches of the trunk lateral line (Sapède et al., 2002). However, the function of *Irx5a* in the primordium is not known (Lecaudey et al., 2005). Furthermore, the lack of reported expression in zebrafish neuromasts contrasts with *Irx5* expression in developing trunk (but not cranial) neuromasts in paddlefish and sterlet, suggesting lineage-specific differences.

Ampullary organ-restricted *Satb2* encodes a homeodomain transcription factor and chromatin-remodeller that is important for craniofacial development, including osteoblast differentiation (reviewed by Huang et al., 2022). Its expression has not been reported in the inner ear or lateral line system. The *Satb2* gene is directly bound by Smad1/5 and upregulated following over-expression of *Bmp4* in cranial neural crest cells, suggesting that *Satb2* is a direct target of the Bmp signalling pathway (Bonilla-Claudio et al., 2012). This raises the possibility that Bmp signalling may be important for ampullary organ development. Indeed, *Bmp4*, *Bmp5*, *Brinp3* (encoding BMP/retinoic acid-inducible neural-specific protein 3) and *Bambi* (encoding a Bmp/activin inhibitor) are present in the ’common’ lateral line organ-enriched gene-set from stage 45 Siberian sturgeon (Wang et al., 2020). *Brinp3* is also in the ampullary organ-enriched gene-set (Wang et al., 2020). Our stage 46 paddlefish lateral line organ-enriched gene-set also includes *Bmp5*, together with genes encoding the dual Bmp/Wnt inhibitors Sostdc1 and Apcdd1 (Modrell et al., 2017a). Thus, the Bmp pathway is a promising target for studies of ampullary organ development.

The other electrosensory-restricted transcription factor genes on the head were *Insm1*, *Mafa*, *Rorc* and *Sp5*. In zebrafish, *Insm1a* is expressed in the migrating posterior lateral line primordium and neuromasts on the trunk (cranial lateral line expression was not reported), and morphants showed defects in primordium migration, proliferation and neuromast formation (He et al., 2017). In the inner ear, transient expression of Insm1 in developing outer hair cells prevents them from transdifferentiating into inner hair cells, by repressing a set of genes usually enriched in early inner hair cells (Wiwatpanit et al., 2018). It is possible, therefore, that Insm1 also acts in developing ampullary organs to repress hair cell-specific genes.

MafA synergises with Neurod1 (and Pdx1) to activate the *insulin* promoter in pancreatic beta-cells (reviewed in Liang et al., 2022). Given the ampullary organ-restricted expression of *Neurod4* in the paddlefish lateral line system (Modrell et al., 2017a), this raises the possibility that MafA could similarly synergise with Neurod4 to activate ampullary organ-specific target genes.

*Rorc* encodes two isoforms of a ligand-dependent transcription factor, RAR-related orphan nuclear receptor gamma (RORγ and RORγt), primarily studied for its roles in regulating Th17 cell differentiation and thus autoimmune and inflammatory diseases (see Fauber and Magnuson, 2014; Meijer et al., 2020; Ladurner et al., 2021). Endogenous ligands for RORγ have not been confirmed, but it responds to sterols including the cholesterol precursor, desmosterol (see Hu et al., 2015; Meijer et al., 2020). Retinoic acid has also been reported to inhibit RORγ activity (Stehlin-Gaon et al., 2003). In the axolotl, ampullary organs were missing and far fewer cranial neuromasts formed after retinoic acid treatment for one hour at late gastrula/early neurula stages (Gibbs and Northcutt, 2004b). However, this most likely reflects an effect on the lateral line placodes themselves, rather than organ formation directly (Gibbs and Northcutt, 2004b). In any case, the mutually exclusive expression of ampullary organ-restricted *Rorc* and cranial neuromast-restricted *Rorb* is particularly intriguing (also see next section).

Finally, *Sp5* encodes a Wnt/β-catenin effector (Kennedy et al., 2016), suggesting that this signalling pathway might be important for ampullary organ development. Indeed, one of the other ampullary organ-restricted genes, *Irx5*, is directly upregulated by Wnt/β-catenin signalling in somatic cells of the gonad (Koth et al., 2020).

Overall, the ampullary organ-restricted cranial expression of these six transcription factor genes, as well as *Neurod4* (Modrell et al., 2017a), provides a starting point for identifying molecular mechanisms that may be important for the formation of electrosensory lateral line organs.

### Five novel mechanosensory lateral line-restricted transcription factor genes

We identified five mechanosensory lateral line-restricted transcription factor genes: the first-such genes reported in electroreceptive vertebrates. Of these, *Hmx2*, *Isl1* and *Rorb* are expressed in zebrafish lateral line placodes and/or neuromasts (Feng and Xu, 2010; Dufourcq et al., 2006; Bertrand et al., 2007). *Hmx2* and *Isl1* both encode homeodomain transcription factors. In zebrafish, *Hmx2* is expressed throughout lateral line placode development, together with the related gene *Hmx3* (Feng and Xu, 2010). Morpholino knockdown experiments suggested a redundant requirement for *Hmx2* and *Hmx3* for cell proliferation in the migrating posterior lateral line primordium, and for normal neuromast formation (Feng and Xu, 2010). Double mutant analysis of *Hmx2* and *Hmx3a* suggested that the loss of neuromasts arises from stalling of the migrating primordium adjacent to the first few somites, hence failure to deposit neuromasts (England et al., 2020). Recent scRNA-seq data from zebrafish also show that *Hmx2* is expressed specifically in anterior-posterior (A/P) support cells in neuromasts (Baek et al., 2022).

We cloned *Isl1* because it promotes a more complete conversion by Atoh1 of mouse cochlear supporting cells to hair cells than does Atoh1 alone (Yamashita et al., 2018). In zebrafish neuromasts, *Isl1* is expressed in multiple support cell types including central support cells (Lush et al., 2019; Baek et al., 2022), which divide symmetrically to form new hair cells after hair cells are ablated (Romero-Carvajal et al., 2015; Lush et al., 2019). In neural crest-derived sensory ganglia, Isl1 is expressed in all neurons and is necessary for nociceptor lineage-specific gene expression, for repressing earlier-acting neurogenic transcription factors - including direct repression of *Neurod4*, which is ampullary organ-specific in the paddlefish lateral line (Modrell et al., 2017a) - and for repressing lineage-inappropriate genes (Sun et al., 2008; Dykes et al., 2011). In the pancreas, Isl1 is a direct transcriptional repressor of *Mafa* (Du et al., 2009), which we identified here as ampullary organ-restricted (see previous section). We hypothesize that, in electroreceptive species, Isl1 may promote a hair cell fate within neuromasts at least in part by repressing an electroreceptor fate, including by repressing *Neurod4* and *Mafa*.

*Rorb*, encoding RAR-related orphan nuclear receptor beta (RORβ), is expressed by supporting cells in adult, regenerating and embryonic neuromasts in zebrafish (Dufourcq et al., 2006; Bertrand et al., 2007). Retinoic acid is a confirmed inhibitory ligand for RORβ (Stehlin-Gaon et al., 2003). However, in sterlet, we only identified *Rorb* expression in cranial neuromasts, suggesting lineage-specific differences. The reciprocal expression of *Rorb* in cranial neuromasts and *Rorc* in ampullary organs (see previous section) suggests that these ligand-dependent transcription factors play specific roles in the development of mechanosensory versus electrosensory organs.

In contrast to *Hmx2*, *Isl1* and *Rorb*, mechanosensory lateral line-restricted *Foxg1* and *Sox8* are not expressed in the developing lateral line system of zebrafish or *Xenopus* (e.g., Dirksen and Jamrich, 1995; Papalopulu and Kintner, 1996; Toresson et al., 1998; Eagleson and Dempewolf, 2002; Duggan et al., 2008; Zhao et al., 2009; O’Donnell et al., 2006; Martik et al., 2019). As zebrafish and *Xenopus* only have a mechanosensory lateral line system (Baker et al., 2013; Baker, 2019), this suggests *Foxg1* and *Sox8* may play specific roles in the developing mechanosensory lateral line system of electroreceptive bony vertebrates, rather than in lateral line primordium or neuromast development *per se*. (In cartilaginous fishes, i.e., sharks, *Sox8* is expressed in ampullary organs as well as neuromasts; this study and Freitas et al., 2006.)

In paddlefish and sterlet, *Foxg1* was expressed in the central zones of lateral line sensory ridges where neuromasts form, though excluded from the central domains of neuromasts where hair cells differentiate. In the mouse olfactory epithelium, Foxg1 maintains a proliferative Sox2^+^ progenitor state (Kawauchi et al., 2009). Similarly, in the inner ear, Foxg1 is expressed by Sox2^+^ hair cell progenitors and supporting cells in sensory epithelia (Kiernan et al., 2005; Dabdoub et al., 2008; Tasdemir-Yilmaz et al., 2021), although it is also expressed by a subset of hair cells (Pauley et al., 2006). *Foxg1* mouse mutants have reduced inner ear sensory epithelia and a shortened cochlea with numerous additional rows of disorganized hair cells (Pauley et al., 2006; Hwang et al., 2009). Conditional knockout of *Foxg1* in supporting cells in the neonatal mouse inner ear resulted in increased numbers of hair cells, potentially by transdifferentiation of supporting cells (Zhang et al., 2019; Zhang et al., 2020). Overall, *Foxg1* is an exciting candidate for further investigation at the functional level.

Recent work on otic placode development in chicken embryos (Buzzi et al., 2022) suggests that another of the chondrostean lateral line mechanosensory-restricted transcription factors we identified, *Sox8*, could play an even earlier role than *Foxg1*. (*Sox8* was expressed in ampullary organs as well as neuromasts in cartilaginous fishes, however, suggesting lineage-specific divergence of expression between cartilaginous and bony vertebrates.) Sox8 in chicken lies upstream of all other transcription factor genes in the otic gene regulatory network, including *Foxg1* (Buzzi et al., 2022). Ectopic expression of *Sox8* in cranial ectoderm drives the formation of ectopic otic vesicles and neurons (Buzzi et al., 2022). In paddlefish, *Sox8* displays a similar expression pattern to *Foxg1* in elongating lateral line primordia and neuromasts. As mentioned, *Sox8* expression has not been reported in the developing lateral line system in either *Xenopus* or zebrafish (O’Donnell et al., 2006; Martik et al., 2019). Given the ’master regulator’ role of *Sox8* in otic placode development (Buzzi et al., 2022), it is possible that *Sox8* lies upstream of *Foxg1* in lateral line primordium development specifically in electroreceptive vertebrates (although it may play a separate, later role in ampullary organ development in cartilaginous fishes).

## Conclusion

The data presented here, taken together with our previous results in paddlefish (Modrell et al., 2011b; Modrell et al., 2011a; Modrell et al., 2017b; Modrell et al., 2017a), show that most transcription factor genes expressed in developing lateral line organs in chondrostean ray-finned fishes, including many that are required for hair cell development, are expressed in both ampullary organs and neuromasts. This supports the hypothesis that the molecular mechanisms underlying electrosensory and mechanosensory lateral line organ development are highly conserved, and that electroreceptors likely evolved as transcriptionally related sister cell types to lateral line hair cells (Modrell et al., 2017a; Baker and Modrell, 2018). Moreover, in addition to ampullary organ-restricted *Neurod4* (Modrell et al., 2017a), we have identified a further eleven transcription factors (six that are electrosensory-restricted on the head; five that are mechanosensory-restricted) that could be involved in the formation of electrosensory versus mechanosensory organs. These are good candidates for functional experiments using CRISPR/Cas9-mediated mutagenesis in sterlet (e.g., Chen et al., 2018; Baloch et al., 2019; Stundl et al., 2022), the next step to further our understanding of the development of these sensory (sister) cell types.

## Materials and Methods

### Embryo collection, staging and fixation

Fertilized sterlet (*Acipenser ruthenus*) eggs were obtained from adults bred at the Research Institute of Fish Culture and Hydrobiology (RIFCH), Faculty of Fisheries and Protection of Waters, University of South Bohemia in České Budějovice, Vodňany, Czech Republic. Sterlet animal husbandry, *in vitro* fertilization and the rearing of embryos and yolk-sac larvae are described in detail in Stundl et al. (2022). Sterlet embryos were staged according to Dettlaff et al. (1993). Animal care was approved by the Ministry of Agriculture of the Czech Republic (MSMT-12550/2016-3), followed the principles of the European Union Harmonized Animal Welfare Act of the Czech Republic, and Principles of Laboratory Animal Care and National Laws 246/1992 “Animal Welfare”, and was conducted in accordance with the Animal Research Committee of RIFCH.

Mississippi paddlefish (*Polyodon spathula*) embryos were purchased from Osage Catfisheries Inc. (Osage Beach, MO, USA) and reared at approximately 22°C in tanks with filtered and recirculating water (pH 7.2 ± 0.7, salinity of 1.0 ± 0.2 ppt). Paddlefish embryos were staged according to Bemis and Grande (1992). Lesser-spotted catshark (*Scyliorhinus canicula*) egg cases were reared in a flow-through seawater system at the Station Biologique de Roscoff, France. Catshark embryos were staged according to Ballard et al. (1993).

Upon reaching desired developmental stages, embryos/larvae of all three species were euthanized via overdose of MS-222 (Sigma-Aldrich). Paddlefish and sterlet embryos/yolk-sac larvae were fixed in modified Carnoy’s fixative (6 volumes 100% ethanol: 3 volumes 37% formaldehyde: 1 volume glacial acetic acid) for 3 hours at room temperature or for 12-24 hours at 4°C, then dehydrated stepwise into ethanol and stored at -20°C. Catshark embryos were fixed overnight at 4°C in 4% paraformaldehyde in phosphate-buffered saline (PBS), washed three times in PBS, dehydrated stepwise into methanol and stored at -20°C.

### Generation of *de novo* transcriptome assemblies from late-larval sterlet heads

Sterlet yolk-sac larvae intended for RNA isolation were preserved in RNAlater (Invitrogen, Thermo Fisher Scientific) and stored at -80°C until processed. Prior to RNA isolation, RNAlater was removed, and heads were manually dissected from sterlet yolk-sac larvae: two at stage 40, two at stage 42, three at stage 45. RNA was then extracted using TRIzol reagent (Invitrogen, Thermo Fisher Scientific) according to the manufacturer’s instructions. RNA concentration was assessed using a Nanodrop N1000 spectrophotometer and integrity using an Agilent 2100 Bioanalyzer (Cambridge Genomic Services, Department of Pathology, University of Cambridge, UK). Samples with an RNA integrity number (RIN) greater than 9 were submitted for next-generation sequencing at The Centre for Applied Genomics, The Hospital for Sick Children, Toronto, Canada. Libraries were prepared using the NEBNExt Ultra Directional RNA library prep kit and sequenced on an Illumina HiSeq 2500, using Illumina v3 chemistry, following the multiplex paired-end protocol (2 x 125 bases).

Reads were subjected to various quality controls, including high-quality read filtering based on the score value given in fastq files (FastQC version 0.10.1; http://www.bioinformatics.babraham.ac.uk/projects/fastqc/), removal of reads containing primer/adaptor sequences and read-length trimming using Trimmomatic-0.30 (Bolger et al., 2014). *De novo* assembly was performed using Velvet version 1.2.10 (Zerbino and Birney, 2008) and Oases version 0.2.08 (Schulz et al., 2012). Velvet was run using different k-mer lengths, k31, k43, k47, k53 and k63 along with other default parameters. Oases was run using the same k-mer range. Results from these assemblies were merged, using Velvet and Oases k-mer of k43. All assemblies were performed on a server with 64 cores and 512 GB of RAM. A second *de novo* assembly was carried out using Trinity version 2.6.6 (Grabherr et al., 2011) using default parameters. This Transcriptome Shotgun Assembly project has been deposited at DDBJ/EMBL/GenBank under the accessions GKLU00000000 (Velvet-Oases assembly) and GKEF00000000 (Trinity assembly). The versions described in this paper are the first versions, GKLU00000000 and GKEF01000000.

### Gene cloning and sequence verification

Total RNA was isolated from embryos using Trizol (Invitrogen, Thermo Fisher Scientific), following the manufacturer’s protocol, and cDNA made using the Superscript III First Strand Synthesis kit (Invitrogen, Thermo Fisher Scientific). To design gene-specific PCR primers or synthetic gene fragments to use as riboprobe templates for *in situ* hybridisation for paddlefish or sterlet, we used the previously published paddlefish transcriptome assembly (NCBI Gene Expression Omnibus accession code GSE92470; Modrell et al., 2017a) or the sterlet transcriptome assemblies reported here (deposited at DDBJ/EMBL/GenBank under the accessions GKLU00000000 and GKEF01000000). Gene-specific primers (Supplementary File 1) were used to amplify cDNA fragments under standard PCR conditions from cDNA and cloned into the pDrive cloning vector (Qiagen) as previously described (Modrell et al., 2011a). Alternatively, synthetic gene fragments based on paddlefish or sterlet transcriptome data, with added M13 forward and reverse primer adaptors, were ordered from Twist Bioscience. To design gene-specific PCR primers for lesser-spotted catshark, we used *S. canicula* RNAseq data, publicly available via the Skatebase website (http://skatebase.org/skateblast-skatebase%e2%80%8b/). Catshark cDNA fragments were cloned into the pGEM-T Easy vector (Promega).

The sterlet and paddlefish riboprobe template sequences were designed prior to the publication of chromosome-level genome assemblies for sterlet (Du et al., 2020) and paddlefish (Cheng et al., 2021). In sterlet, roughly 70% of ohnologues (i.e., gene paralogs resulting from an independent whole-genome duplication in the sterlet lineage) proved to have been retained (Du et al., 2020). The paddlefish underwent an independent species-specific whole-genome duplication relatively recently (Cheng et al., 2021). Both ohnologues have been retained for all genes described here except sterlet *Foxi2*, sterlet *Rorc* and paddlefish *Sox10*. Supplementary File 1 includes each riboprobe’s percentage match with each ohnologue, obtained using the National Center for Biotechnology Information (NCBI) Basic Local Alignment Search Tool (BLAST; https://blast.ncbi.nlm.nih.gov/Blast.cgi; McGinnis and Madden, 2004) by performing a nucleotide BLAST search against the respective genome assemblies. The percentage match with the ’targeted’ ohnologue ranged from 97.5-100% for sterlet (mean ± s.d. 99.6% ± 0.55; n=39) and from 98.7-100% for paddlefish (mean ± s.d. 99.6% ± 0.40; n=9). The percentage match with the second ohnologue was also high, ranging from 87.4-100% for sterlet (mean ± s.d. 97.0% ± 2.71; n=37) and from 90.7-99.0% for paddlefish (mean ± s.d. 94.4% ± 2.57, n=8) (Supplementary File 1), suggesting that our riboprobes most likely also target transcripts from the second ohnologue, where present. Indeed, three of our paddlefish riboprobes (*Irx5*, *Lhx8* and *Sox2*) also worked well in sterlet; the percentage match with the top-match sterlet ohnologue ranged from 93.5% to 96.7% (Supplementary File 1).

GenBank accession numbers for sterlet (*A. ruthenus*), paddlefish (*P. spathula*) and catshark (*S. canicula*) cDNA fragments, synthetic gene fragments or predicted transcripts from the sterlet or paddlefish genomes are given in Supplementary File 1, as are the nucleotide ranges targeted by our riboprobes. The sterlet *Rorc* sequence was absent from the reference genome assembly (ASM1064508v2; Du et al., 2020), but present in the Vertebrate Genomes Project chromosome-level sterlet assembly (fAciRut3.2 paternal haplotype; GCA_902713435.1), which is available via Rapid Ensembl (https://rapid.ensembl.org/).

Individual clones were verified by sequencing (Department of Biochemistry Sequencing Facility, University of Cambridge, UK, or Genewiz, Azenta Life Sciences, UK). Sequence identity was checked using the NCBI BLAST tool. Sequences whose identity was still inconclusive following a general BLAST search were checked against the sterlet reference genome (ASM1064508v2; Du et al., 2020) or paddlefish reference genome (ASM1765450v1; Cheng et al., 2021) using BLAST. However, we note here that this approach did not result in conclusive identification of our *Insm* family gene transcripts and a *Klf* gene transcript. We thus performed phylogenetic analysis of these gene families using predicted protein sequences from reference genome assemblies of a range of species of deuterostomes. The accession numbers for these sequences are listed in Supplementary File 2. The sequences were aligned using MAFFT (Katoh and Standley, 2013) and trimmed using TrimAL (Capella-Gutiérrez et al., 2009) before using IQ-TREE2 (Minh et al., 2020) with Model Finder (Kalyaanamoorthy et al., 2017) for phylogenetic tree inference and bootstrap analysis. Trees were then visualised using TreeGraph 2 (Stöver and Müller, 2010). Our phylogenetic analysis of *Insm* family genes revealed that the *Insm2* ohnologues in the reference sterlet genome (Du et al., 2020) have been mis-annotated as *Insm1* and *Insm1-like*, while in the reference paddlefish genome (Cheng et al., 2021), one of the *Insm2* ohnologues has been mis-annotated as *Insm1a-like* (Supplementary Figure S4; Supplementary File 1). Similarly, our phylogenetic analysis of *Klf* family genes revealed that one of the *Klf17* ohnologues in the reference sterlet genome (Du et al., 2020) has been mis-annotated as *Klf4* (Supplementary Figures S5 and S6; Supplementary File 1).

### *In situ* hybridization and immunohistochemistry

Digoxigenin-labelled antisense riboprobes were synthesized from cloned cDNA fragments (Supplementary file 1) using T7 or SP6 polymerases (Promega) and digoxigenin-labelled dUTPs (Roche). Alternatively, synthetic gene fragments (Twist Bioscience) with added M13 forward and reverse primer adaptors were PCR-amplified under standard conditions using the M13 forward primer, and the M13 reverse primer containing an overhang with the SP6 polymerase promoter. The PCR product was then used as a template for riboprobe synthesis by *in vitro* transcription using SP6 polymerase and digoxigenin-labelled dUTPs (Roche). Each riboprobe was tested a minimum of two times, using at least three embryos per stage.

Wholemount in situ hybridization (ISH) was performed as previously described (Modrell et al., 2011a). In some cases, sterlet and paddlefish yolk-sac larvae were processed into pre-hybridization buffer as described (Modrell et al., 2011a), then stored at -20°C for up to a month in this solution before continuing the protocol. For weaker riboprobes, overnight incubations at 4°C in MABT (0.1 M maleic acid, 150 mM NaCl, 0.1% Tween-20, pH 7.5) and/or NTMT (100 mM NaCl, 100 mM Tris, pH 9.5, 50 mM MgCl2, 0.1% Tween-20) were added prior to the colour reaction, to increase the signal to background staining ratio.

Wholemount immunostaining was performed as previously described (Metscher and Müller, 2011). When using sterlet embryos or yolk-sac larvae that had not already been subject to ISH, bleaching and proteinase K treatment were performed prior to immunostaining, as described for ISH (Modrell et al., 2011a). A primary antibody against Sox2 (rabbit monoclonal, ab92494; Abcam) was used at 1:200 and a horseradish peroxidase-conjugated goat anti-rabbit antibody (Jackson ImmunoResearch) at 1:300. For the histochemical reaction, the metallographic peroxidase substrate EnzMet kit (Nanoprobes) was used according to the manufacturer’s instructions.

For skinmounts after wholemount ISH and/or immunostaining, skin samples were dissected using forceps and microcapillary needles and mounted on Superfrost Plus slides (VWR) using Fluoroshield mounting medium with DAPI (Sigma-Aldrich).

For ISH on sections, embryos were embedded in paraffin wax and sectioned at 10 μm as previously described (O’Neill et al., 2007). ISH on sections was performed as previously described (O’Neill et al., 2007; Miller et al., 2017) except that slides were not treated with proteinase K prior to hybridization and BMP Purple (Roche) was used for the colour reaction.

### Imaging and image processing

Wholemount embryos and larvae were positioned in a slit in an agar-coated Petri dish with PBS and imaged using a Leica MZFLIII dissecting microscope equipped with a MicroPublisher 5.0 RTV camera (QImaging) or a MicroPublisher 6 color CCD camera (Teledyne Photometrics). Skinmounts and sections were imaged using a Zeiss AxioSkop 2 microscope equipped with a Retiga 2000R camera and RGB pancake (QImaging) or a MicroPublisher 6 color CCD camera (Teledyne Photometrics). Images were acquired using QCapture Pro 6.0 or 7.0 software (QImaging) or Ocular software (Teledyne Photometrics). For most whole-mount embryos and larvae, as well as skinmounts, a stack of images was taken by manually focusing through the sample, then focus stacking was performed using Helicon Focus software (Helicon Soft Limited). Images were processed in Adobe Photoshop (Adobe Systems Inc.).

## Data availability statement

The Transcriptome Shotgun Assembly project has been deposited at DDBJ/EMBL/GenBank under the accessions GKLU00000000 and GKEF01000000. The versions described in this paper are the first versions, GKLU00000000 and GKEF01000000. The publication and associated supplementary figures include representative example images of embryos from each experiment. Additional raw data underlying this publication consist of further images of these and other embryos from each experiment. Public sharing of these images is not cost-efficient, but they are available from the corresponding author upon reasonable request.

## Ethics statement

Sterlet animal work was reviewed and approved by The Animal Research Committee of Research Institute of Fish Culture and Hydrobiology, Faculty of Fisheries and Protection of Waters, University of South Bohemia in České Budějovice, Vodňany, Czech Republic and Ministry of Agriculture of the Czech Republic (MSMT-12550/2016-3).

## Author contributions

CB conceived and designed the project with input from MSM, MM and AG, and wrote the manuscript together with MM. MM performed most of the experiments, prepared all the manuscript figures and made a significant contribution to the writing of the manuscript. MSM, AG, AC and IF performed some experiments. RL undertook the transcriptome assembly with support from GM. MP and DG were instrumental in enabling all work with sterlet embryos. All authors read and commented on the manuscript.

## Rights Retention Statement

This work was funded by grants from the Biotechnology and Biological Sciences Research Council (BB/F00818X/1 and BB/P001947/1). For the purpose of open access, the author has applied a Creative Commons Attribution (CC BY) licence to any Author Accepted Manuscript version arising.

## Acknowledgments

Thanks to Marek Rodina and Martin Kahanec for their help with sterlet spawns, and Michaela Vazačová for her help with embryo incubation and fixation. Thanks to Tatjana Piotrowski and her lab at the Stowers Institute for Medical Research (Kansas City, MO, USA) and Steve and Pete Kahrs and the Kahrs family (Osage Catfisheries, Inc.) for hosting MSM during paddlefish spawning seasons. Thanks to Christine Hirschberger and Rolf Ericsson for their help with some of the *in situ* hybridization rounds. Thanks to Nathanael Walker-Hale for advice on phylogenetic analysis.

## References

1. Baek, S., Tran, N.T.T., Diaz, D.C., Tsai, Y.-Y., Acedo, J.N., Lush, M.E., Piotrowski, T., 2022. Single-cell transcriptome analysis reveals three sequential phases of gene expression during zebrafish sensory hair cell regeneration. Dev. Cell 57, 799–819.

2. Baker, C.V.H., 2019. The development and evolution of lateral line electroreceptors: insights from comparative molecular approaches., in: B.A. Carlson, J.A. Sisneros, A.N. Popper, R.R. Fay (Eds.), Electroreception: Fundamental Insights from Comparative Approaches. Springer, Cham, pp. 25–62.

3. Baker, C.V.H., Modrell, M.S., 2018. Insights into electroreceptor development and evolution from molecular comparisons with hair cells. Integr. Comp. Biol. 58, 329–340.

4. Baker, C.V.H., Modrell, M.S., Gillis, J.A., 2013. The evolution and development of vertebrate lateral line electroreceptors. J. Exp. Biol. 216, 2515–2522.

5. Ballard, W.W., Mellinger, J., Lechenault, H., 1993. A series of normal stages for development of *Scyliorhinus canicula*, the lesser spotted dogfish (*Chondrichthyes*: *Scylorhinidae*). J. Exp. Zool. 267, 318–336.

6. Baloch, A.R., Franěk, R., Tichopád, T., Fučíková, M., Rodina, M., Pšenička, M., 2019. *Dnd1* knockout in sturgeons by CRISPR/Cas9 generates germ cell free host for surrogate production. Animals (Basel) 9, 174.

7. Bardot, E.S., Valdes, V.J., Zhang, J., Perdigoto, C.N., Nicolis, S., Hearn, S.A., Silva, J.M., Ezhkova, E., 2013. Polycomb subunits Ezh1 and Ezh2 regulate the Merkel cell differentiation program in skin stem cells. EMBO J. 32, 1990–2000.

8. Bellono, N.W., Leitch, D.B., Julius, D., 2017. Molecular basis of ancestral vertebrate electroreception. Nature 543, 391–396.

9. Bellono, N.W., Leitch, D.B., Julius, D., 2018. Molecular tuning of electroreception in sharks and skates. Nature 558, 122–126.

10. Bemis, W.E., Grande, L., 1992. Early development of the actinopterygian head. I. External development and staging of the paddlefish Polyodon spathula. J. Morphol. 213, 47–83.

11. Bennett, M.V.L., Obara, S., 1986. Ionic mechanisms and pharmacology of electroreceptors, in: T.H. Bullock, W. Heiligenberg (Eds.), Electroreception. Wiley, New York, pp. 157–181.

12. Bertrand, S., Thisse, B., Tavares, R., Sachs, L., Chaumot, A., Bardet, P.L., Escrivà, H., Duffraisse, M., Marchand, O., Safi, R., Thisse, C., Laudet, V., 2007. Unexpected novel relational links uncovered by extensive developmental profiling of nuclear receptor expression. PLoS Genet. 3, e188.

13. Bodznick, D., Montgomery, J.C., 2005. The physiology of low-frequency electrosensory systems, in: T.H. Bullock, C.D. Hopkins, A.N. Popper, R.R. Fay (Eds.), Electroreception. Springer, New York, pp. 132–153.

14. Bolger, A.M., Lohse, M., Usadel, B., 2014. Trimmomatic: a flexible trimmer for Illumina sequence data. Bioinformatics 30, 2114–2120.

15. Bonilla-Claudio, M., Wang, J., Bai, Y., Klysik, E., Selever, J., Martin, J.F., 2012. Bmp signaling regulates a dose-dependent transcriptional program to control facial skeletal development. Development 139, 709–719.

16. Bosse, A., Stoykova, A., Nieselt-Struwe, K., Chowdhury, K., Copeland, N.G., Jenkins, N.A., Gruss, P., 2000. Identification of a novel mouse Iroquois homeobox gene, *Irx5*, and chromosomal localisation of all members of the mouse Iroquois gene family. Dev. Dyn. 218, 160–174.

17. Brown, T.L., Horton, E.C., Craig, E.W., Goo, C.E.A., Black, E.C., Hewitt, M.N., Yee, N.G., Fan, E.T., Raible, D.W., Rasmussen, J.P., 2023. Dermal appendage-dependent patterning of zebrafish *atoh1a*+ Merkel cells. eLife 12, e85800.

18. Butts, T., Modrell, M.S., Baker, C.V.H., Wingate, R.J.T., 2014. The evolution of the vertebrate cerebellum: absence of a proliferative external granule layer in a non-teleost ray-finned fish. Evol. Dev. 16, 92–100.

19. Buzzi, A.L., Chen, J., Thiery, A., Delile, J., Streit, A., 2022. Sox8 remodels the cranial ectoderm to generate the ear. Proc. Natl. Acad. Sci. U.S.A. 119, e2118938119.

20. Camacho, S., Ostos, M.D.V., Llorente, J.I., Sanz, A., García, M., Domezain, A., Carmona, R., 2007. Structural characteristics and development of ampullary organs in *Acipenser naccarii*. Anat. Rec. (Hoboken) 290, 1178–1189.

21. Capella-Gutiérrez, S., Silla-Martínez, J.M., Gabaldón, T., 2009. trimAl: a tool for automated alignment trimming in large-scale phylogenetic analyses. Bioinformatics 25, 1972–1973.

22. Caprara, G.A., Peng, A.W., 2022. Mechanotransduction in mammalian sensory hair cells. Mol. Cell. Neurosci. 120, 103706.

23. Cardeña-Núñez, S., Sánchez-Guardado, L.Ó., Corral-San-Miguel, R., Rodríguez-Gallardo, L., Marín, F., Puelles, L., Aroca, P., Hidalgo-Sánchez, M., 2016. Expression patterns of Irx genes in the developing chick inner ear. Brain Struct. Funct. 222, 2071–2092.

24. Chagnaud, B.P., Wilkens, L.A., Hofmann, M., 2021. The ampullary electrosensory system –a paddlefish case study, in: B. Fritzsch (Ed.), The Senses: A Comprehensive Reference. Elsevier, pp. 215–227.

25. Chen, J., Wang, W., Tian, Z., Dong, Y., Dong, T., Zhu, H., Zhu, Z., Hu, H., Hu, W., 2018. Efficient gene transfer and gene editing in sterlet. Front. Genet. 9, 117.

26. Chen, Y., Gu, Y., Li, Y., Li, G.L., Chai, R., Li, W., Li, H., 2021. Generation of mature and functional hair cells by co-expression of *Gfi1*, *Pou4f3*, and *Atoh1* in the postnatal mouse cochlea. Cell Rep. 35, 109016.

27. Cheng, C.W., Chow, R.L., Lebel, M., Sakuma, R., Cheung, H.O.-L., Thanabalasingham, V., Zhang, X., Bruneau, B.G., Birch, D.G., Hui, C.C., McInnes, R.R., Cheng, S.H., 2005. The *Iroquois* homeobox gene, *Irx5*, is required for retinal cone bipolar cell development. Dev. Biol. 287, 48–60.

28. Cheng, P., Huang, Y., Lv, Y., Du, H., Ruan, Z., Li, C., Ye, H., Zhang, H., Wu, J., Wang, C., Ruan, R., Li, Y., Bian, C., You, X., Shi, C., Han, K., Xu, J., Shi, Q., Wei, Q., 2021. The American paddlefish genome provides novel insights into chromosomal evolution and bone mineralization in early vertebrates. Mol. Biol. Evol. 38, 1595–1607.

29. Crampton, W.G.R., 2019. Electroreception, electrogenesis and electric signal evolution. J. Fish Biol. 95, 92–134.

30. Dabdoub, A., Puligilla, C., Jones, J.M., Fritzsch, B., Cheah, K.S., Pevny, L.H., Kelley, M.W., 2008. Sox2 signaling in prosensory domain specification and subsequent hair cell differentiation in the developing cochlea. Proc. Natl. Acad. Sci. U.S.A. 105, 18396–18401.

31. Deprez, M., Zaragosi, L.E., Truchi, M., Becavin, C., Ruiz García, S., Arguel, M.J., Plaisant, M., Magnone, V., Lebrigand, K., Abelanet, S., Brau, F., Paquet, A., Pe’er, D., Marquette, C.H., Leroy, S., Barbry, P., 2020. A single-cell atlas of the human healthy airways. Am. J. Respir. Crit. Care Med. 202, 1636–1645.

32. Dettlaff, T.A., Ginsburg, A.S., Schmalhausen, O.I., 1993. Sturgeon Fishes: Developmental Biology and Aquaculture. Springer-Verlag, Berlin.

33. Dirksen, M.-L., Jamrich, M., 1995. Differential expression of *fork head* genes during early *Xenopus* and zebrafish development. Dev. Genet. 17, 107–116.

34. Du, A., Hunter, C.S., Murray, J., Noble, D., Cai, C.-L., Evans, S.M., Stein, R., May, C.L., 2009. Islet-1 is required for the maturation, proliferation, and survival of the endocrine pancreas. Diabetes 58, 2059–2069.

35. Du, K., Stöck, M., Kneitz, S., Klopp, C., Woltering, J.M., Adolfi, M.C., Feron, R., Prokopov, D., Makunin, A., Kichigin, I., Schmidt, C., Fischer, P., Kuhl, H., Wuertz, S., Gessner, J., Kloas, W., Cabau, C., Iampietro, C., Parrinello, H., Tomlinson, C., Journot, L., Postlethwait, J.H., Braasch, I., Trifonov, V., Warren, W.C., Meyer, A., Guiguen, Y., Schartl, M., 2020. The sterlet sturgeon genome sequence and the mechanisms of segmental rediploidization. Nat. Ecol. Evol. 4, 841–852.

36. Dufourcq, P., Roussigné, M., Blader, P., Rosa, F., Peyrieras, N., Vriz, S., 2006. Mechano-sensory organ regeneration in adults: the zebrafish lateral line as a model. Mol. Cell. Neurosci. 33, 180–187.

37. Duggan, C.D., Demaria, S., Baudhuin, A., Stafford, D., Ngai, J., 2008. Foxg1 is required for development of the vertebrate olfactory system. J. Neurosci. 28, 5229–5239.

38. Dykes, I.M., Tempest, L., Lee, S.-I., Turner, E.E., 2011. Brn3a and Islet1 act epistatically to regulate the gene expression program of sensory differentiation. J. Neurosci. 31, 9789–9799.

39. Eagleson, G.W., Dempewolf, R.D., 2002. The role of the anterior neural ridge and *Fgf-8* in early forebrain patterning and regionalization in *Xenopus laevis*. Comp. Biochem. Physiol. B Biochem. Mol. Biol. 132, 179–189.

40. England, S.J., Cerda, G.A., Kowalchuk, A., Sorice, T., Grieb, G., Lewis, K.E., 2020. Hmx3a has essential functions in zebrafish spinal cord, ear and lateral line development. Genetics 216, 1153–1185.

41. Fauber, B.P., Magnuson, S., 2014. Modulators of the nuclear receptor retinoic acid receptor-related orphan receptor-γ (RORγ or RORc). J. Med. Chem. 57, 5871–5892.

42. Feng, Y., Xu, Q., 2010. Pivotal role of hmx2 and hmx3 in zebrafish inner ear and lateral line development. Dev. Biol. 339, 507–518.

43. Freitas, R., Zhang, G., Albert, J.S., Evans, D.H., Cohn, M.J., 2006. Developmental origin of shark electrosensory organs. Evol. Dev. 8, 74–80.

44. Gibbs, M.A., Northcutt, R.G., 2004a. Development of the lateral line system in the shovelnose sturgeon. Brain Behav. Evol. 64, 70–84.

45. Gibbs, M.A., Northcutt, R.G., 2004b. Retinoic acid repatterns axolotl lateral line receptors. Int. J. Dev. Biol. 48, 63–66.

46. Gillis, J.A., Modrell, M.S., Northcutt, R.G., Catania, K.C., Luer, C.A., Baker, C.V.H., 2012. Electrosensory ampullary organs are derived from lateral line placodes in cartilaginous fishes. Development 139, 3142–3146.

47. Gnedeva, K., Hudspeth, A.J., 2015. SoxC transcription factors are essential for the development of the inner ear. Proc. Natl. Acad. Sci. U.S.A. 112, 14066–14071.

48. Grabherr, M.G., Haas, B.J., Yassour, M., Levin, J.Z., Thompson, D.A., Amit, I., Adiconis, X., Fan, L., Raychowdhury, R., Zeng, Q., Chen, Z., Mauceli, E., Hacohen, N., Gnirke, A., Rhind, N., di Palma, F., Birren, B.W., Nusbaum, C., Lindblad-Toh, K., Friedman, N., Regev, A., 2011. Full-length transcriptome assembly from RNA-Seq data without a reference genome. Nat. Biotechnol. 29, 644–652.

49. Haber, A.L., Biton, M., Rogel, N., Herbst, R.H., Shekhar, K., Smillie, C., Burgin, G., Delorey, T.M., Howitt, M.R., Katz, Y., Tirosh, I., Beyaz, S., Dionne, D., Zhang, M., Raychowdhury, R., Garrett, W.S., Rozenblatt-Rosen, O., Shi, H.N., Yilmaz, O., Xavier, R.J., Regev, A., 2017. A single-cell survey of the small intestinal epithelium. Nature 551, 333–339.

51. He, Y., Lu, X., Qian, F., Liu, D., Chai, R., Li, H., 2017. *Insm1a* is required for zebrafish posterior lateral line development. Front. Mol. Neurosci. 10, 241.

52. Hesse, K., Vaupel, K., Kurt, S., Buettner, R., Kirfel, J., Moser, M., 2011. AP-2δ is a crucial transcriptional regulator of the posterior midbrain. PLoS ONE 6, e23483.

53. Hoffman, B.U., Baba, Y., Griffith, T.N., Mosharov, E.V., Woo, S.H., Roybal, D.D., Karsenty, G., Patapoutian, A., Sulzer, D., Lumpkin, E.A., 2018. Merkel cells activate sensory neural pathways through adrenergic synapses. Neuron 100, 1401–1413.e6.

54. Houweling, A.C., Dildrop, R., Peters, T., Mummenhoff, J., Moorman, A.F.M., Rüther, U., Christoffels, V.M., 2001. Gene and cluster-specific expression of the *Iroquois* family members during mouse development. Mech. Dev. 107, 169–174.

55. Hu, X., Wang, Y., Hao, L.-Y., Liu, X., Lesch, C.A., Sanchez, B.M., Wendling, J.M., Morgan, R.W., Aicher, T.D., Carter, L.L., Toogood, P.L., Glick, G.D., 2015. Sterol metabolism controls T(H)17 differentiation by generating endogenous RORγ agonists. Nat. Chem. Biol. 11, 141–147.

56. Huang, X., Chen, Q., Luo, W., Pakvasa, M., Zhang, Y., Zheng, L., Li, S., Yang, Z., Zeng, H., Liang, F., Zhang, F., Hu, D.A., Qin, K.H., Wang, E.J., Qin, D.S., Reid, R.R., He, T.-C., Athiviraham, A., El Dafrawy, M., Zhang, H., 2022. SATB2: A versatile transcriptional regulator of craniofacial and skeleton development, neurogenesis and tumorigenesis, and its applications in regenerative medicine. Genes Dis. 9, 95–107.

57. Hwang, C.H., Simeone, A., Lai, E., Wu, D.K., 2009. *Foxg1* is required for proper separation and formation of sensory cristae during inner ear development. Dev. Dyn. 238, 2725–2734.

58. Iyer, A.A., Groves, A.K., 2021. Transcription factor reprogramming in the inner ear: turning on cell fate switches to regenerate sensory hair cells. Front. Cell. Neurosci. 15, 660748.

59. Iyer, A.A., Hosamani, I., Nguyen, J.D., Cai, T., Singh, S., McGovern, M.M., Beyer, L., Zhang, H., Jen, H.I., Yousaf, R., Birol, O., Sun, J.J., Ray, R.S., Raphael, Y., Segil, N., Groves, A.K., 2022. Cellular reprogramming with ATOH1, GFI1, and POU4F3 implicate epigenetic changes and cell-cell signaling as obstacles to hair cell regeneration in mature mammals. eLife 11, e79712.

60. Jan, T.A., Eltawil, Y., Ling, A.H., Chen, L., Ellwanger, D.C., Heller, S., Cheng, A.G., 2021. Spatiotemporal dynamics of inner ear sensory and non-sensory cells revealed by single-cell transcriptomics. Cell Rep. 36, 109358.

61. Jen, H.-I., Singh, S., Tao, L., Maunsell, H.R., Segil, N., Groves, A.K., 2022. GFI1 regulates hair cell differentiation by acting as an off-DNA transcriptional co-activator of ATOH1, and a DNA-binding repressor. Sci. Rep. 12, 7793.

62. Jessen, K.R., Mirsky, R., 2019. Schwann cell precursors; multipotent glial cells in embryonic nerves. Front. Mol. Neurosci. 12, 69.

63. Jørgensen, J.M., 2005. Morphology of electroreceptive sensory organs, in: T.H. Bullock, C.D. Hopkins, A.N. Popper, R.R. Fay (Eds.), Electroreception. Springer, New York, pp. 47–67.

64. Kalyaanamoorthy, S., Minh, B.Q., Wong, T.K.F., von Haeseler, A., Jermiin, L.S., 2017. ModelFinder: fast model selection for accurate phylogenetic estimates. Nat. Methods 14, 587–589.

65. Katoh, K., Standley, D.M., 2013. MAFFT multiple sequence alignment software version 7: improvements in performance and usability. Mol. Biol. Evol. 30, 772–780.

66. Kawauchi, S., Santos, R., Kim, J., Hollenbeck, P.L.W., Murray, R.C., Calof, A.L., 2009. The role of *Foxg1* in the development of neural stem cells of the olfactory epithelium. Ann. N. Y. Acad. Sci. 1170, 21–27.

67. Kennedy, M.W., Chalamalasetty, R.B., Thomas, S., Garriock, R.J., Jailwala, P., Yamaguchi, T.P., 2016. Sp5 and Sp8 recruit β-catenin and Tcf1-Lef1 to select enhancers to activate Wnt target gene transcription. Proc. Natl. Acad. Sci. U.S.A. 113, 3545–3550.

68. Kiernan, A.E., Pelling, A.L., Leung, K.K., Tang, A.S., Bell, D.M., Tease, C., Lovell-Badge, R., Steel, K.P., Cheah, K.S., 2005. *Sox2* is required for sensory organ development in the mammalian inner ear. Nature 434, 1031–1035.

69. Kotas, M.E., O’Leary, C.E., Locksley, R.M., 2023. Tuft cells: context- and tissue-specific programming for a conserved cell lineage. Annu. Rev. Pathol. 18, 311–335.

70. Koth, M.L., Garcia-Moreno, S.A., Novak, A., Holthusen, K.A., Kothandapani, A., Jiang, K., Taketo, M.M., Nicol, B., Yao, H.H.-C., Futtner, C.R., Maatouk, D.M., Jorgensen, J.S., 2020. Canonical Wnt/β-catenin activity and differential epigenetic marks direct sexually dimorphic regulation of *Irx3* and *Irx5* in developing mouse gonads. Development 147, dev183814.

71. Kotkamp, K., Mössner, R., Allen, A., Onichtchouk, D., Driever, W., 2014. A Pou5f1/Oct4 dependent Klf2a, Klf2b, and Klf17 regulatory sub-network contributes to EVL and ectoderm development during zebrafish embryogenesis. Dev. Biol. 385, 433–447.

72. Ladurner, A., Schwarz, P.F., Dirsch, V.M., 2021. Natural products as modulators of retinoic acid receptor-related orphan receptors (RORs). Nat. Prod. Rep. 38, 757–781.

73. Lecaudey, V., Anselme, I., Dildrop, R., Rüther, U., Schneider-Maunoury, S., 2005. Expression of the zebrafish Iroquois genes during early nervous system formation and patterning. J. Comp. Neurol. 492, 289–302.

74. Leitch, D.B., Julius, D., 2019. Electrosensory transduction: comparisons across structure, afferent response properties, and cellular physiology., in: B.A. Carlson, J.A. Sisneros, A.N. Popper, R.R. Fay (Eds.), Electroreception: Fundamental Insights from Comparative Approaches. Springer, Cham, pp. 63–90.

75. Lesko, M.H., Driskell, R.R., Kretzschmar, K., Goldie, S.J., Watt, F.M., 2013. Sox2 modulates the function of two distinct cell lineages in mouse skin. Dev. Biol. 382, 15–26.

76. Li, X., Gaillard, F., Monckton, E.A., Glubrecht, D.D., Persad, A.R., Moser, M., Sauvé, Y., Godbout, R., 2016. Loss of AP-2delta reduces retinal ganglion cell numbers and axonal projections to the superior colliculus. Mol. Brain 9, 62.

77. Liang, J., Chirikjian, M., Pajvani, U.B., Bartolomé, A., 2022. MafA Regulation in β-Cells: From Transcriptional to Post-Translational Mechanisms. Biomolecules 12, 535.

78. Lorenzen, S.M., Duggan, A., Osipovich, A.B., Magnuson, M.A., García-Añoveros, J., 2015. Insm1 promotes neurogenic proliferation in delaminated otic progenitors. Mech. Dev. 138 Pt 3, 233–245.

79. Lu, X., Sipe, C.W., 2016. Developmental regulation of planar cell polarity and hair-bundle morphogenesis in auditory hair cells: lessons from human and mouse genetics. WIREs Dev. Biol. 5, 85–101.

80. Lush, M.E., Diaz, D.C., Koenecke, N., Baek, S., Boldt, H., St Peter, M.K., Gaitan-Escudero, T., Romero-Carvajal, A., Busch-Nentwich, E.M., Perera, A.G., Hall, K.E., Peak, A., Haug, J.S., Piotrowski, T., 2019. scRNA-Seq reveals distinct stem cell populations that drive hair cell regeneration after loss of Fgf and Notch signaling. eLife 8, e44431.

81. Mak, A.C.Y., Szeto, I.Y.Y., Fritzsch, B., Cheah, K.S.E., 2009. Differential and overlapping expression pattern of SOX2 and SOX9 in inner ear development. Gene Expr. Patterns 9, 444–453.

82. Martik, M.L., Gandhi, S., Uy, B.R., Gillis, J.A., Green, S.A., Simoes-Costa, M., Bronner, M.E., 2019. Evolution of the new head by gradual acquisition of neural crest regulatory circuits. Nature 574, 675–678.

83. Matern, M.S., Milon, B., Lipford, E.L., McMurray, M., Ogawa, Y., Tkaczuk, A., Song, Y., Elkon, R., Hertzano, R., 2020. GFI1 functions to repress neuronal gene expression in the developing inner ear hair cells. Development 147, dev186015.

84. McGinnis, S., Madden, T.L., 2004. BLAST: at the core of a powerful and diverse set of sequence analysis tools. Nucleic Acids Res. 32, W20–5.

85. Meijer, F.A., Doveston, R.G., de Vries, R.M.J.M., Vos, G.M., Vos, A.A.A., Leysen, S., Scheepstra, M., Ottmann, C., Milroy, L.-G., Brunsveld, L., 2020. Ligand-based design of allosteric retinoic acid receptor-related orphan receptor γt (RORγt) inverse agonists. J. Med. Chem. 63, 241–259.

86. Metscher, B.D., Müller, G.B., 2011. MicroCT for molecular imaging: quantitative visualization of complete three-dimensional distributions of gene products in embryonic limbs. Dev. Dyn. 240, 2301–2308.

87. Miller, S.R., Perera, S.N., Baker, C.V.H., 2017. Constitutively active Notch1 converts cranial neural crest-derived frontonasal mesenchyme to perivascular cells *in vivo*. Biol. Open 6, 317–325.

88. Minh, B.Q., Schmidt, H.A., Chernomor, O., Schrempf, D., Woodhams, M.D., von Haeseler, A., Lanfear, R., 2020. IQ-TREE 2: New models and efficient methods for phylogenetic inference in the genomic era. Mol. Biol. Evol. 37, 1530–1534.

89. Modrell, M.S., Baker, C.V.H., 2012. Evolution of electrosensory ampullary organs: conservation of *Eya4* expression during lateral line development in jawed vertebrates. Evol. Dev. 14, 277–285.

90. Modrell, M.S., Bemis, W.E., Northcutt, R.G., Davis, M.C., Baker, C.V.H., 2011a. Electrosensory ampullary organs are derived from lateral line placodes in bony fishes. Nat. Commun. 2, 496.

91. Modrell, M.S., Buckley, D., Baker, C.V.H., 2011b. Molecular analysis of neurogenic placode development in a basal ray-finned fish. genesis 49, 278–294.

92. Modrell, M.S., Lyne, M., Carr, A.R., Zakon, H.H., Buckley, D., Campbell, A.S., Davis, M.C., Micklem, G., Baker, C.V.H., 2017a. Insights into electrosensory organ development, physiology and evolution from a lateral line-enriched transcriptome. eLife 6, e24197.

93. Modrell, M.S., Tidswell, O.R.A., Baker, C.V.H., 2017b. Notch and Fgf signaling during electrosensory versus mechanosensory lateral line organ development in a non-teleost ray-finned fish. Dev. Biol. 431, 48–58.

94. Mogdans, J., 2019. Sensory ecology of the fish lateral-line system: Morphological and physiological adaptations for the perception of hydrodynamic stimuli. J. Fish Biol. 95, 53–72.

95. Moser, T., Grabner, C.P., Schmitz, F., 2020. Sensory processing at ribbon synapses in the retina and the cochlea. Physiol. Rev. 100, 103–144.

96. Norris, H.W., Hughes, S.P., 1920. The spiracular sense-organ in elasmobranchs, ganoids and dipnoans. Anat. Rec. 18, 205–209.

97. Northcutt, R.G., 1997. Evolution of gnathostome lateral line ontogenies. Brain Behav. Evol. 50, 25–37.

98. Northcutt, R.G., Brändle, K., Fritzsch, B., 1995. Electroreceptors and mechanosensory lateral line organs arise from single placodes in axolotls. Dev. Biol. 168, 358–373.

99. Ó Maoiléidigh, D., Ricci, A.J., 2019. A bundle of mechanisms: inner-ear hair-cell mechanotransduction. Trends Neurosci. 42, 221–236.

100. O’Donnell, M., Hong, C.-S., Huang, X., Delnicki, R.J., Saint-Jeannet, J.-P., 2006. Functional analysis of Sox8 during neural crest development in *Xenopus*. Development 133, 3817–3826.

101. O’Neill, P., McCole, R.B., Baker, C.V.H., 2007. A molecular analysis of neurogenic placode and cranial sensory ganglion development in the shark, *Scyliorhinus canicula*. Dev. Biol. 304, 156–181.

102. Pangrsic, T., Singer, J.H., Koschak, A., 2018. Voltage-gated calcium channels: key players in sensory coding in the retina and the inner ear. Physiol Rev. 98, 2063–2096.

103. Papalopulu, N., Kintner, C., 1996. A posteriorising factor, retinoic acid, reveals that anteroposterior patterning controls the timing of neuronal differentiation in *Xenopus* neuroectoderm. Development 122, 3409–3418.

104. Pauley, S., Lai, E., Fritzsch, B., 2006. *Foxg1* is required for morphogenesis and histogenesis of the mammalian inner ear. Dev. Dyn. 235, 2470–2482.

105. Perdigoto, C.N., Bardot, E.S., Valdes, V.J., Santoriello, F.J., Ezhkova, E., 2014. Embryonic maturation of epidermal Merkel cells is controlled by a redundant transcription factor network. Development 141, 4690–4696.

106. Piotrowski, T., Baker, C.V.H., 2014. The development of lateral line placodes: Taking a broader view. Dev. Biol. 389, 68–81.

107. Reilly, M.B., Cros, C., Varol, E., Yemini, E., Hobert, O., 2020. Unique homeobox codes delineate all the neuron classes of *C. elegans*. Nature 584, 595–601.

108. Roccio, M., Senn, P., Heller, S., 2020. Novel insights into inner ear development and regeneration for targeted hearing loss therapies. Hear. Res. 397, 107859.

109. Romero-Carvajal, A., Navajas Acedo, J., Jiang, L., Kozlovskaja-Gumbriene, A., Alexander, R., Li, H., Piotrowski, T., 2015. Regeneration of sensory hair cells requires localized interactions between the Notch and Wnt pathways. Dev. Cell 34, 267–282.

110. Saito, T., Psenicka, M., 2015. Novel technique for visualizing primordial germ cells in sturgeons (*Acipenser ruthenus*, *A. gueldenstaedtii*, *A. baerii*, and *Huso huso*). Biol. Reprod. 93, 96.

111. Sapède, D., Gompel, N., Dambly-Chaudière, C., Ghysen, A., 2002. Cell migration in the postembryonic development of the fish lateral line. Development 129, 605–615.

112. Schulz, M.H., Zerbino, D.R., Vingron, M., Birney, E., 2012. Oases: robust *de novo* RNA-seq assembly across the dynamic range of expression levels. Bioinformatics 28, 1086–1092.

113. Stehlin-Gaon, C., Willmann, D., Zeyer, D., Sanglier, S., Van Dorsselaer, A., Renaud, J.-P., Moras, D., Schüle, R., 2003. All-*trans* retinoic acid is a ligand for the orphan nuclear receptor ROR beta. Nat. Struct. Biol. 10, 820–825.

114. Stöver, B.C., Müller, K.F., 2010. TreeGraph 2: combining and visualizing evidence from different phylogenetic analyses. BMC Bioinformatics 11, 7.

115. Stundl, J., Martik, M.L., Chen, D., Raja, D.A., Franěk, R., Pospisilova, A., Pšenička, M., Metscher, B.D., Braasch, I., Haitina, T., Cerny, R., Ahlberg, P.E., Bronner, M.E., 2023. Ancient vertebrate dermal armor evolved from trunk neural crest. Proc. Natl. Acad. Sci. U.S.A. 120, e2221120120.

116. Stundl, J., Pospisilova, A., Matějková, T., Psenicka, M., Bronner, M.E., Cerny, R., 2020. Migratory patterns and evolutionary plasticity of cranial neural crest cells in ray-finned fishes. Dev. Biol. 467, 14–29.

117. Stundl, J., Soukup, V., Franěk, R., Pospisilova, A., Psutkova, V., Pšenička, M., Cerny, R., Bronner, M.E., Medeiros, D.M., Jandzik, D., 2022. Efficient CRISPR mutagenesis in sturgeon demonstrates its utility in large, slow-maturing vertebrates. Front. Cell Dev. Biol. 10, 750833.

118. Sun, Y., Dykes, I.M., Liang, X., Eng, S.R., Evans, S.M., Turner, E.E., 2008. A central role for Islet1 in sensory neuron development linking sensory and spinal gene regulatory programs. Nat. Neurosci. 11, 1283–1293.

119. Tasdemir-Yilmaz, O.E., Druckenbrod, N.R., Olukoya, O.O., Dong, W., Yung, A.R., Bastille, I., Pazyra-Murphy, M.F., Sitko, A.A., Hale, E.B., Vigneau, S., Gimelbrant, A.A., Kharchenko, P.V., Goodrich, L.V., Segal, R.A., 2021. Diversity of developing peripheral glia revealed by single-cell RNA sequencing. Dev. Cell 56, 2516–2535.

120. Toresson, H., Martinez-Barbera, J.P., Bardsley, A., Caubit, X., Krauss, S., 1998. Conservation of *BF-1* expression in amphioxus and zebrafish suggests evolutionary ancestry of anterior cell types that contribute to the vertebrate telencephalon. Dev. Genes Evol. 208, 431–439.

121. Undurraga, C.A., Gou, Y., Sandoval, P.C., Nuñez, V.A., Allende, M.L., Riley, B.B., Hernández, P.P., Sarrazin, A.F., 2019. Sox2 and Sox3 are essential for development and regeneration of the zebrafish lateral line. bioRxiv 856088; doi: 10.1101/856088.

122. Vidal, B., Gulez, B., Cao, W.X., Leyva-Díaz, E., Reilly, M.B., Tekieli, T., Hobert, O., 2022. The enteric nervous system of the *C. elegans* pharynx is specified by the Sine oculis-like homeobox gene *ceh-34*. eLife 11, e76003.

123. von Bartheld, C.S., Giannessi, F., 2011. The paratympanic organ: a barometer and altimeter in the middle ear of birds? J. Exp. Zool. B Mol. Dev. Evol. 316, 402–408.

124. Wang, J., Lu, C., Zhao, Y., Tang, Z., Song, J., Fan, C., 2020. Transcriptome profiles of sturgeon lateral line electroreceptor and mechanoreceptor during regeneration. BMC Genomics 21, 875.

125. Wang, X., Llamas, J., Trecek, T., Shi, T., Tao, L., Makmura, W., Crump, J.G., Segil, N., Gnedeva, K., 2023. SoxC transcription factors shape the epigenetic landscape to establish competence for sensory differentiation in the mammalian organ of Corti. Proc. Natl. Acad. Sci. U.S.A. 120, e2301301120.

126. Webb, J.F., 2021. Morphology of the mechanosensory lateral line system of fishes, in: B. Fritzsch (Ed.), The Senses: A Comprehensive Reference. Elsevier, pp. 29–46.

127. Wiwatpanit, T., Lorenzen, S.M., Cantú, J.A., Foo, C.Z., Hogan, A.K., Márquez, F., Clancy, J.C., Schipma, M.J., Cheatham, M.A., Duggan, A., García-Añoveros, J., 2018. Trans-differentiation of outer hair cells into inner hair cells in the absence of INSM1. Nature 563, 691–695.

128. Yamashita, T., Zheng, F., Finkelstein, D., Kellard, Z., Robert, C., Rosencrance, C.D., Sugino, K., Easton, J., Gawad, C., Zuo, J., 2018. High-resolution transcriptional dissection of *in vivo* Atoh1-mediated hair cell conversion in mature cochleae identifies Isl1 as a co-reprogramming factor. PLoS Genet. 14, e1007552.

129. Zeiske, E., Kasumyan, A., Bartsch, P., Hansen, A., 2003. Early development of the olfactory organ in sturgeons of the genus *Acipenser*: a comparative and electron microscopic study. Anat. Embryol. (Berl.) 206, 357–372.

130. Zerbino, D.R., Birney, E., 2008. Velvet: algorithms for de novo short read assembly using de Bruijn graphs. Genome Res. 18, 821–829.

131. Zhang, S., Zhang, Y., Dong, Y., Guo, L., Zhang, Z., Shao, B., Qi, J., Zhou, H., Zhu, W., Yan, X., Hong, G., Zhang, L., Zhang, X., Tang, M., Zhao, C., Gao, X., Chai, R., 2019. Knockdown of *Foxg1* in supporting cells increases the trans-differentiation of supporting cells into hair cells in the neonatal mouse cochlea. Cell. Mol. Life Sci. 77, 1401–1419.

132. Zhang, Y., Zhang, S., Zhang, Z., Dong, Y., Ma, X., Qiang, R., Chen, Y., Gao, X., Zhao, C., Chen, F., He, S., Chai, R., 2020. Knockdown of *Foxg1* in Sox9+ supporting cells increases the trans-differentiation of supporting cells into hair cells in the neonatal mouse utricle. Aging (Albany NY) 12, 19834–19851.

133. Zhao, X.-F., Suh, C.S., Prat, C.R., Ellingsen, S., Fjose, A., 2009. Distinct expression of two *foxg1* paralogues in zebrafish. Gene Expr. Patterns 9, 266–272.

