## Supplementary Figure_S1-S5 for "Identification of multiple transcription factor genes potentially involved in the development of electrosensory versus mechanosensory lateral line organs"

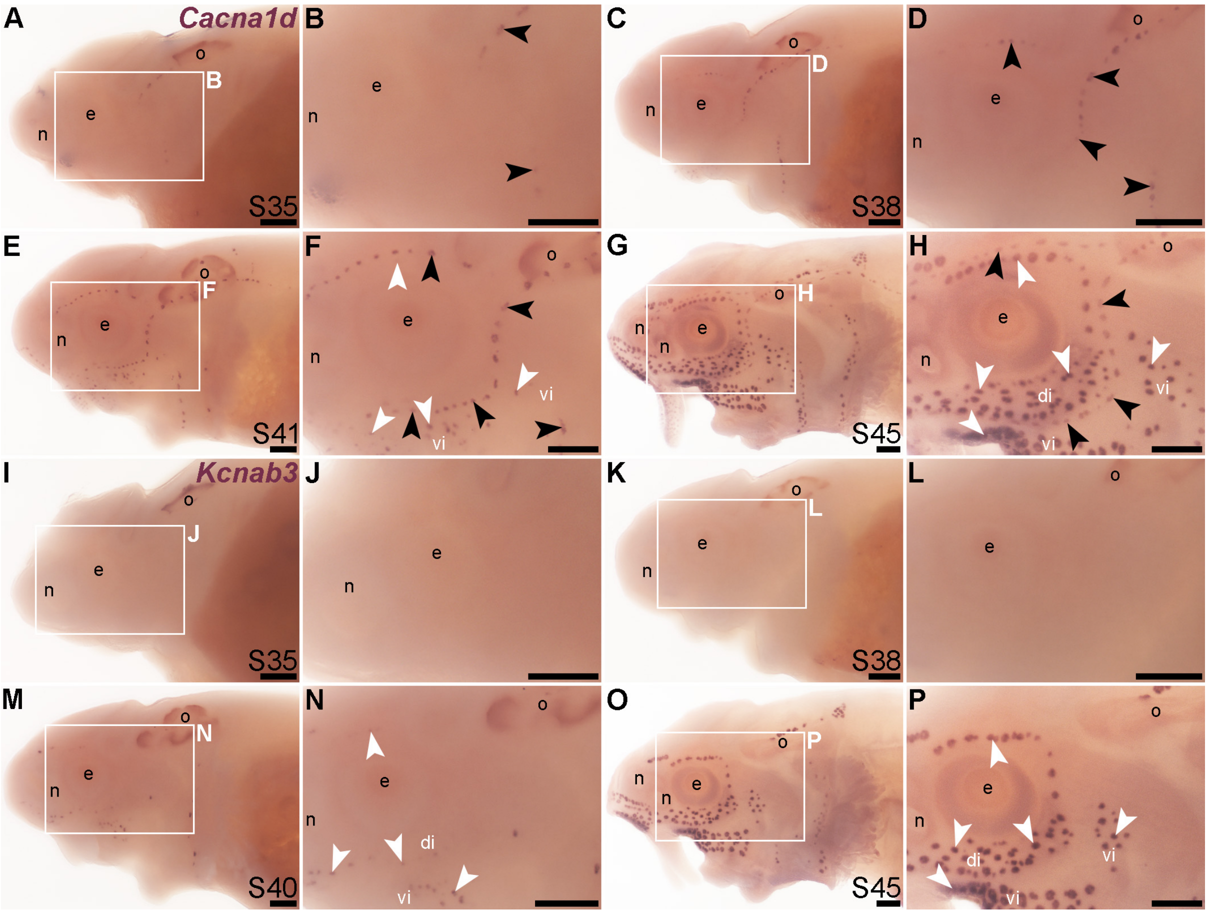


**Supplementary Figure S1. Time-course of neuromast and ampullary organ differentiation in sterlet.** *In situ* hybridization at selected stages in sterlet, from stage 35 (the stage before hatching occurs, at stage 36) to stage 45, the onset of independent feeding. Black arrowheads indicate examples of neuromasts; white arrowheads indicate examples of ampullary organs. (**A-H**) Expression of *Cacna1d*, encoding the pore-forming alpha subunit of the voltage-gated calcium channel $Ca_v1.3$, reveals differentiated hair cells in a few neuromasts already at stage 35 in the otic line, near the otic vesicle (A,B), with numbers increasing and further neuromast line divisions becoming apparent at stage 38 (C,D). Some differentiated electroreceptors are detected already at stage 41 (E,F). All neuromast lines and ampullary organ fields are developed by stage 45 (G,H). *Cacna1d* is also weakly expressed in taste buds, most clearly on the barbels (E,G). (**I-P**) Expression of electroreceptor-specific *Kcnab3* (encoding an accessory subunit for a voltage-gated $K^+$ channel, $K_v\beta3$) is not seen at stages 35 (I,J) or 38 (K,L), but some differentiated electroreceptors are present by stage 40 (M,N). All ampullary organ fields are developed at stage 45 (O,P). Abbreviations: di, dorsal infraorbital ampullary organ field; e, eye; n, naris; o, otic vesicle; S, stage; vi, ventral infraorbital ampullary organ field. Scale bar: 200 μm.

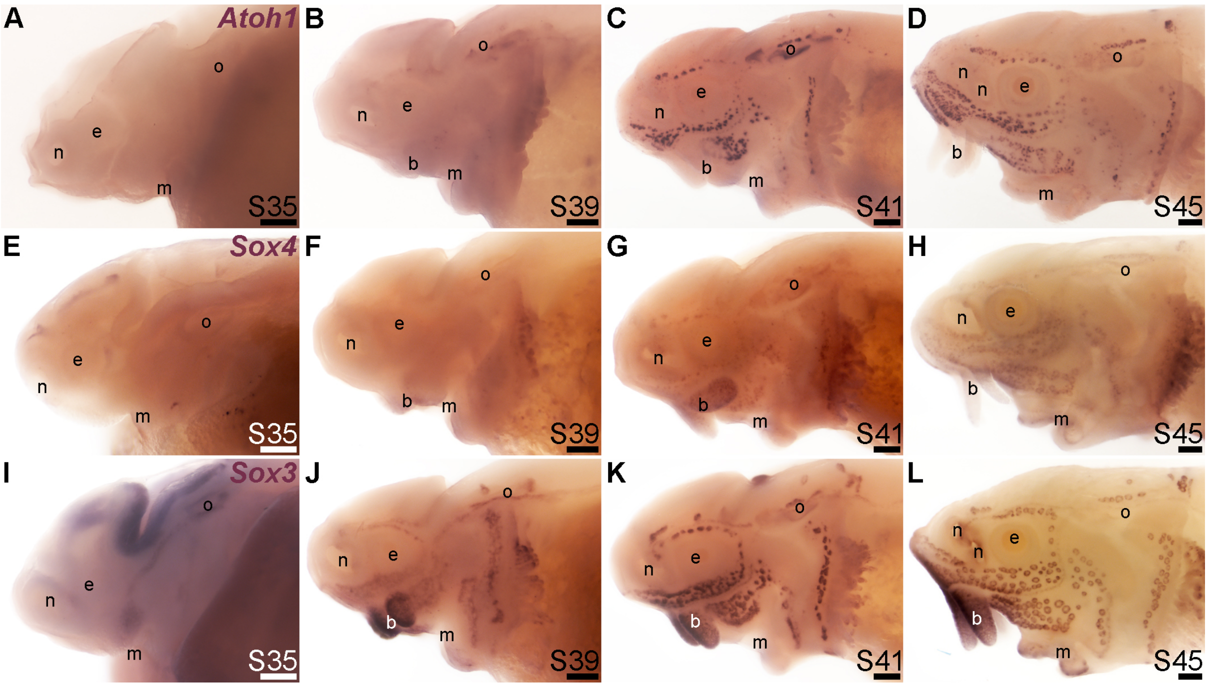


**Supplementary Figure S2. Time-course of *Atoh1, Sox4* and *Sox3* expression in the developing sterlet lateral line system.** *In situ* hybridization at selected stages in sterlet, from stage 35 (the stage before hatching occurs, at stage 36) to stage 45, the onset of independent feeding. (**A-D**) *Atoh1* expression is not seen at stage 35 (A). Expression in developing lateral line organs is weak at stage 39 (B) and well defined by stage 41 (C). At this stage, as well as stage 45 (D), *Atoh1* is expressed more strongly in ampullary organs than in neuromasts. (**E-H**) *Sox4* expression is not seen at stages 35 (E) or 39 (F), but can be detected in ampullary organs and more weakly in neuromasts at stage 41 (G). The same pattern is maintained at stage 45 (H). (**I-L**) *Sox3* is expressed in sensory ridges at stage 35 (**I**), as well as in the otic vesicle, brain, and developing barbels. It is detected in ampullary organs on the operculum at stage 39 (J), and is well defined in all ampullary organ fields by stage 41, with weaker expression in neuromast lines (K). The same pattern is preserved at stage 45 (L). Abbreviations: b, barbels; e, eye; m, mouth; n, naris; o, otic vesicle; S, stage. Scale bar: 200 μm.

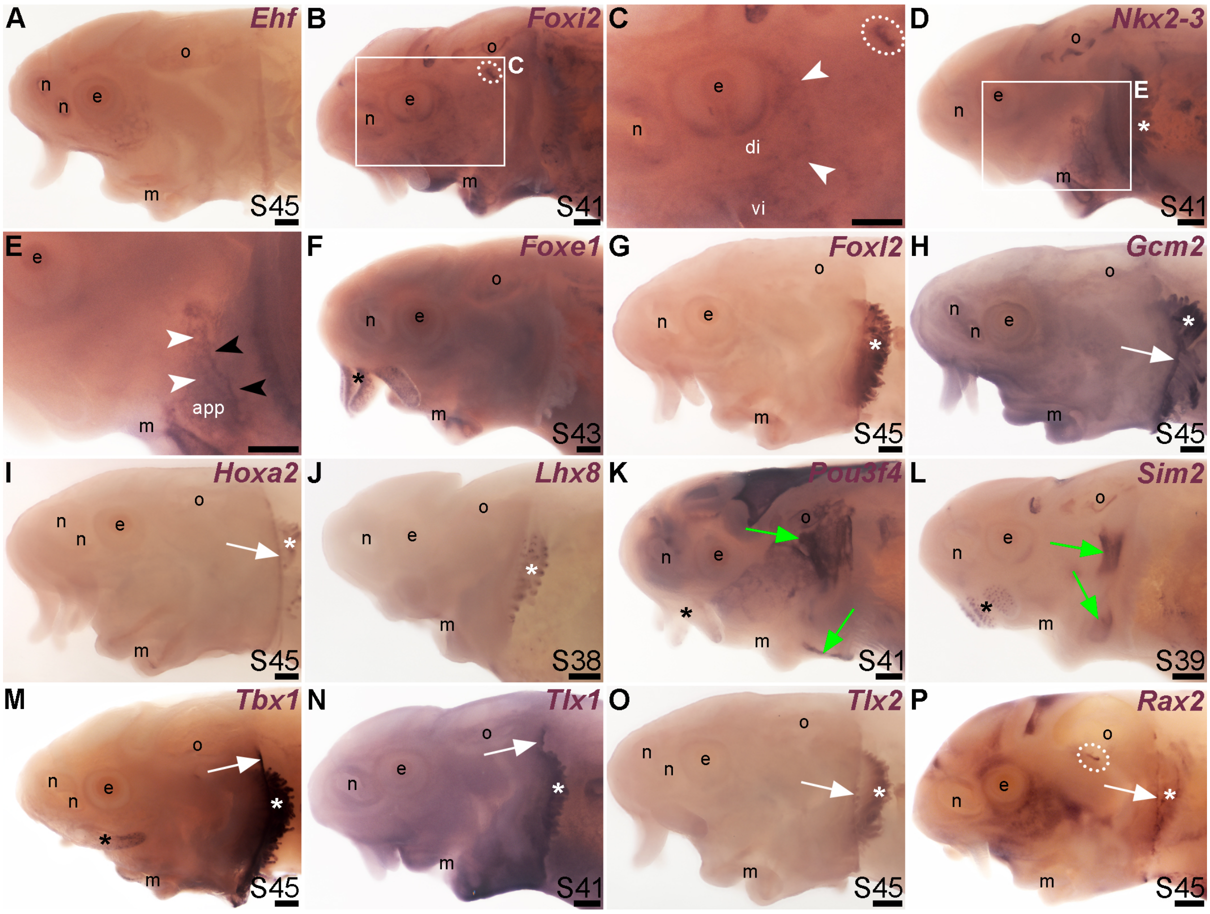


**Supplementary Figure S3. Transcription factor genes not expressed in sterlet lateral line organs.** *In situ* hybridization at selected stages in sterlet, showing genes cloned from the paddlefish stage 46 lateral line organ-enriched gene set (Modrell et al., 2017a) that were not expressed in lateral line organs. (**A**) *Ehf* expression is detected in the skin immediately around ampullary organs and neuromasts of the infraorbital line at stage 45. (**B,C**) *Foxi2* is expressed in the skin immediately around ampullary organs and neuromasts of the infraorbital line at stage 41. *Foxi2* expression is also detected in the brain and otic vesicle, around the mouth, and around the spiracular opening (circled with white dots). (The *Foxi2* sequence was annotated in the paddlefish dataset as *forkhead box protein I1c-like*, but the cloned sterlet sequence matches a gene annotated as *Foxi2* in the sterlet reference genome; Modrell et al., 2017a; Du et al., 2020.) (**D,E**) *Nkx2-3* is expressed in the skin immediately around ampullary organs and neuromasts of the preopercular line at stage 41. (**F**) *Foxe1* expression is seen in taste buds on the barbels (black asterisk) and in and around the mouth at stage 43. (**G**) *Foxl2* expression is seen in gill filaments (white asterisk) at stage 45. (**H**) *Gcm2* is expressed in gill filaments (white asterisk) and at the edge of the operculum (white arrow) at stage 45. (**I**) *Hoxa2* expression is detected in gill filaments (white asterisk) and at the edge of the operculum (white arrow) at stage 45. (**J**) *Lhx8* expression is detected (using the paddlefish riboprobe) in gill filaments (white asterisk) at stage 38. (**K**) *Pou3f4* expression is seen in head mesenchyme at stage 41, as well as some muscles. Green arrows indicate the regions corresponding to the hyohyoideus muscle (top arrow) or interhyoideus muscle (bottom arrow). The barbels (black asterisk) show internal expression at the tip. (Dark staining in the hindbrain is trapping.) (**L**) *Sim2* expression is detected in taste buds on barbels (black asterisk) as well as some muscles at stage 39 (possibly the developing hyohyoideus muscle, green arrow). (**M**) Expression of *Tbx1* (originally unassigned locus 3098; Modrell et al., 2017a) is seen in gill filaments (white asterisk), barbels (black asterisk), and the edge of the operculum (white arrow) at stage 45. (**N**) *Tlx1* expression is seen in gill filaments (white asterisk) and at the edge of the operculum (white arrow) at stage 41. (**O**) *Tlx2* expression is detected in gill filaments (white asterisk) and at the edge of the operculum (white arrow) at stage 45. (**P**) *Rax2* expression is seen around the spiracular opening (circled with white dots) and at the edge of the operculum (white arrow), as well as in the branchial arches and head mesenchyme, at stage 45. Dark staining in the hindbrain is trapping; staining in the otic capsule may also be trapping. (The *Rax2* sequence was annotated in the paddlefish dataset as *visual system homeobox 2-like*, but the cloned sterlet sequence matches a gene annotated as *Rax2* in the sterlet reference genome; Modrell et al., 2017a; Du et al., 2020.) Abbreviations: app, anterior preopercular ampullary organ field; di, dorsal infraorbital ampullary organ field; e, eye; m, mouth; n, naris; o, otic vesicle; S, stage; t, trapping of the colour reaction product in brain ventricles; vi, ventral infraorbital ampullary organ field. Scale bar: 200 µm.

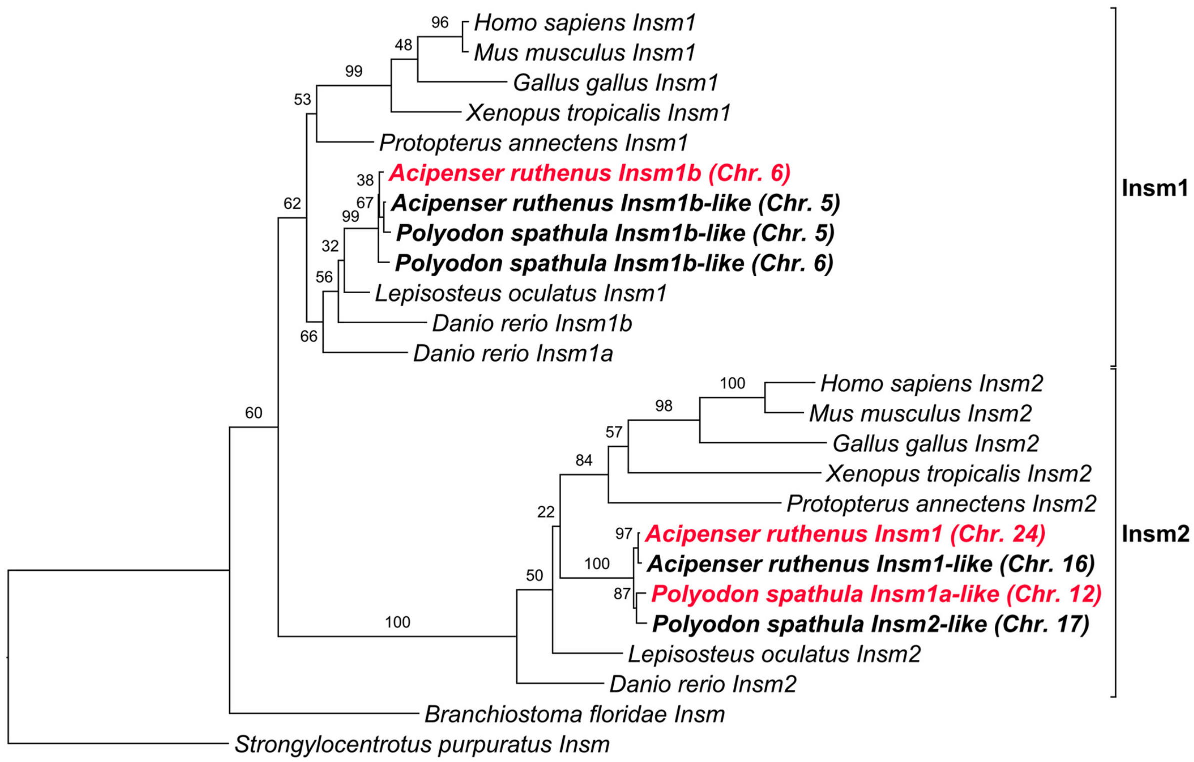


**Supplementary Figure S4. Phylogenetic analysis reveals the identity of *Insm* genes in paddlefish and sterlet.** Phylogenetic tree generated using available amino acid sequences from selected deuterostome reference genomes. Maximum bootstrap support for the *Insm2* clade indicates that both sterlet *Insm2* ohnologues (on chromosomes 16 and 24) and one paddlefish *Insm2* ohnologue (on chromosome 12) have been mis-annotated as *Insm1* in the respective reference genomes (sterlet ASM1064508v2; paddlefish ASM1765450v1; Du et al., 2020; Cheng et al., 2021). The original annotations from the reference genomes and the chromosomal location of each paddlefish and sterlet gene (in parentheses) are provided. The top-match ohnologues for the *Insm1* and *Insm2* riboprobes used in this study are highlighted in red. Numbers at nodes represent bootstrap values obtained from 100 replicates. GenBank accession numbers for the sequences used are provided in Supplementary File 2.

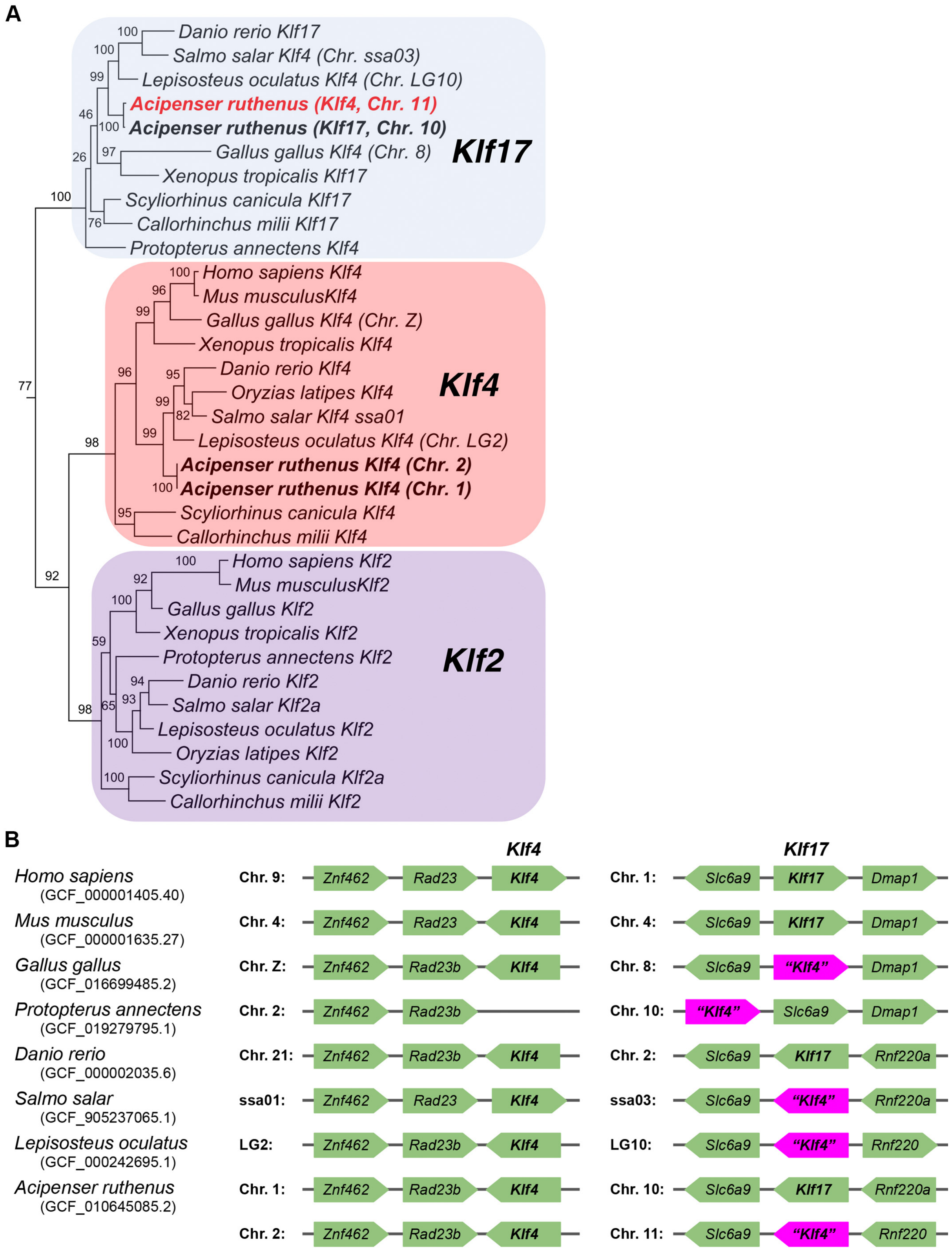


**Supplementary Figure S5. Phylogenetic analysis and synteny relationships confirm the identity of *Klf* genes in sterlet.** (**A**) Selected portion of a phylogenetic tree generated using available amino acid sequences from selected deuterostome reference genomes points to incorrect annotation of several *Klf4/Klf17* genes in selected vertebrate reference genomes within the clade containing all Klf2, Klf4 and Klf17 sequences except mouse and human Klf17 (see Supplementary Figure S6 for the complete tree). This was previously noted by Kotkamp et al. (2014), who also argued that the divergence of mammalian *Klf17* genes is due to rapid evolution of this gene specifically in mammals.

We suggest that all sequences in the mixed Klf4/Klf17 clade, sister to the clade containing all the remaining Klf4 sequences plus Klf2 sequences, should be annotated as Klf17, preserving a well-supported monophyly for each of the *Klf2*, *Klf4* and *Klf17* genes, with the exception of mammalian *Klf17*. Hence, the gene currently annotated as *Klf4* on chromosome 11 in the sterlet reference genome (ASM1064508v2; Du et al., 2020) should be annotated as *Klf17*. The chromosomal locations of the sterlet genes and the mis-annotated gene are provided in parentheses. All sterlet sequences are highlighted in bold. The top-match sterlet ohnologue for the *Klf17* riboprobe used in this study (mis-annotated in the genome as *Klf4*) is highlighted in red. Numbers at nodes represent bootstrap values obtained from 100 replicates. GenBank accession numbers for the sequences used are provided in Supplementary File 2. (**B**) Synteny analysis of genes neighbouring *Klf4* and *Klf17* in selected vertebrate reference genomes confirms mis-annotation of the *Klf17* gene in the reference genomes of chicken *(Gallus gallus)*, African lungfish *(Protopterus annectens)*, Atlantic salmon *(Salmo salar)*, spotted gar *(Lepisosteus oculatus)* and sterlet *(Acipenser ruthenus)*. NCBI RefSeq accession numbers for the relevant reference genomes are shown in parentheses. Chromosomal location is specified next to each synteny schematic. Mis-annotated genes are highlighted in magenta, with the original (incorrect) annotation in quotation marks.
