## Supplementary figures and images for "Identification of multiple transcription factor genes potentially involved in the development of electrosensory versus mechanosensory lateral line organs"

### Supplementary Figure_S6

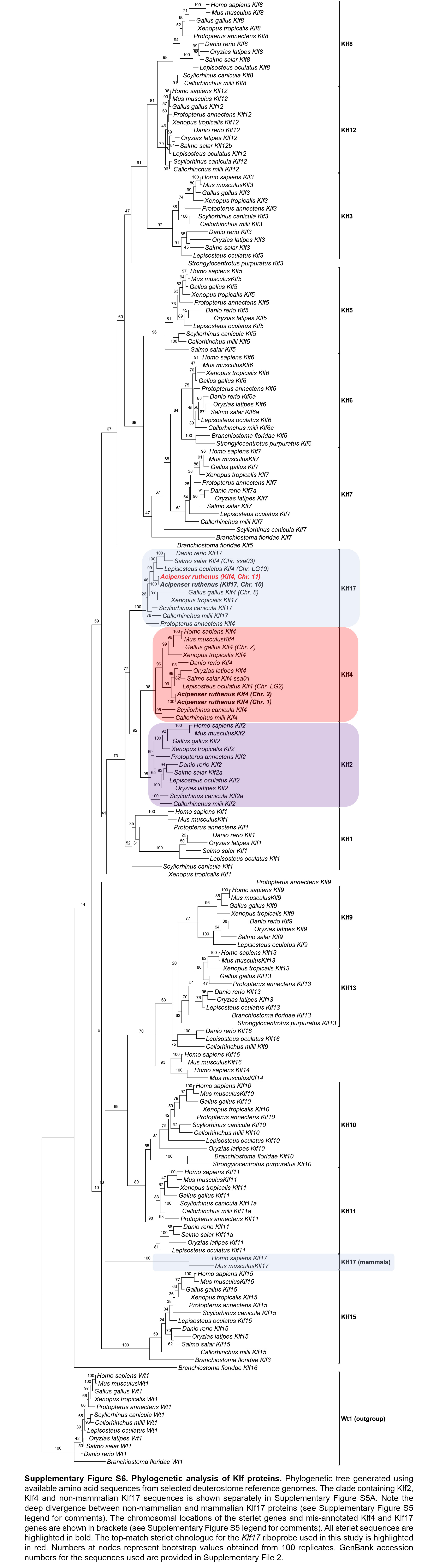
